# Tissue geometry and mechanochemical feedback initiate rotational migration in *Drosophila*

**DOI:** 10.1101/2025.09.03.674060

**Authors:** Sierra Schwabach, Sreejith Santhosh, Audrey Miller Williams, Maureen Cetera, Mattia Serra, Sally Horne-Badovinac

## Abstract

Collective migration of epithelial cells drives diverse tissue remodeling processes. In many cases, a free tissue edge polarizes the cells to promote directed motion, but how edge-free or closed epithelia initiate migration remains unclear. Here, we show that the rotational migration of follicular epithelial cells in the *Drosophila* egg chamber is a self-organizing process. Combining experiments and theoretical modeling, we identify a positive feedback loop in which the mechanosensitive behavior of the atypical cadherin Fat2 synergizes with the rigid-body dynamics of the egg chamber to both initiate and sustain rotation. Mechanical constraints arising from cell–cell interactions and tissue geometry further align this motion around the egg chamber’s anterior–posterior axis. Our findings reveal a biophysical mechanism — combining Fat2-mediated velocity–polarity alignment, rigid-body dynamics, and tissue geometry — by which a closed epithelial tissue self-organizes into persistent, large-scale rotational migration *in vivo*, expanding current flocking theories.

## Introduction

Collective migration of epithelial cells plays central roles in morphogenesis, intestinal turnover, wound repair, and metastasis [1–4]. To migrate collectively, each epithelial cell must establish a protrusive leading edge and a contractile trailing edge. These individual cell migratory behaviors must also be globally oriented across the tissue plane, allowing the entire epithelium to move in a directed manner [5]. When an epithelium has a free edge, as in a wound healing scenario, this physical asymmetry can work alone or with other external cues to polarize the tissue for directed movement [6–8]. There are cases, however, in which epithelial migration occurs in the absence of external cues. Epithelial cells plated on circular micropatterns can spontaneously break chiral symmetry and undergo rotational migration in either a clockwise or counterclockwise direction [9–13]. Cultured epithelial spheres, [14–21], epithelia grown on cylindrical surfaces, [22], and the spherical alveoli of human mammary organoids also rotate persistently [23]. These examples suggest that the ability to self-organize for rotational migration may be a fundamental property of epithelia that adopt distinct geometries [24], but the mechanisms by which rotation initiates are unknown.

Here, we study a rotational migration that occurs in the *Drosophila* egg chamber. An egg chamber is an ovarian follicle that consists of a central cluster of germ cells, surrounded by an epithelium of follicle cells [25, 26]. Egg chambers form in a stem cell compartment called the germarium and then progress through 14 developmental stages as they develop into an egg (Fig. 1A). During the early stages of egg chamber development, the follicle cells collectively migrate along the basement membrane (BM) extracellular matrix that encapsulates the tissue [27, 28].

**Figure 1.**
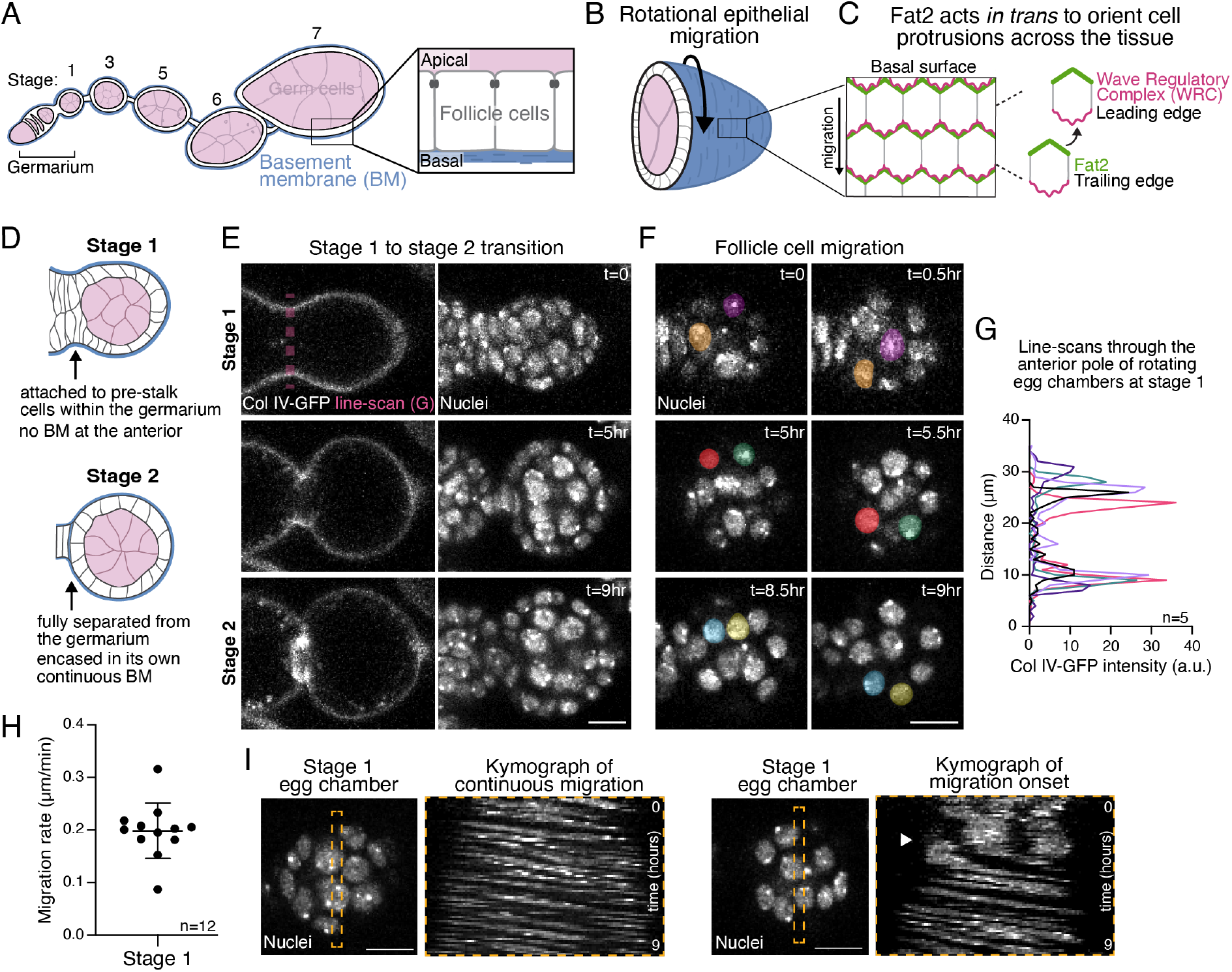
Egg chamber rotation begins within the germarium. **A)** Illustration of a transverse section through a developmental array of egg chambers (ovariole). The boxed region highlights the organization of the follicular epithelium. **B)** Illustration of rotational epithelial migration (arrow), which is driven by follicle cell crawling along the stationary basement membrane. **C)** Illustration of the basal epithelial surface showing how Fat2 acts *in trans* from the trailing edge of each cell to stabilize WRC activity at the leading edge of the cell behind. **D)** Illustrations of transverse sections through stage 1 and 2 egg chambers, highlighting the morphological differences between them. **E)** Movie stills of a transverse section through one egg chamber that capture the transition from stage 1 to 2. Pink dashed line shows the location of the line-scans used to generate the plot in G. Nuclei marked with SpyDNA. **F)** Movie stills focused on the follicular epithelium of the same egg chamber as in E. Two different cells are pseudocolored in each row of images to show cell movement over 30 minutes at each of the three time points. **G)** Line-scans of Col IV-GFP through the anterior-most pole of the follicular epithelium for 5 different rotating egg chambers at stage 1. The bimodal distribution indicates that the BM has not yet formed over the egg chamber’s anterior. **H)** Quantification of follicle cell migration rates for egg chambers at stage 1. Bar represents mean *±* SD. **I)** Movie stills of stage 1 egg chambers with corresponding kymographs that show either constant migration or the onset of migration (arrowhead). Scale bars, 10µm.

This causes the entire egg chamber to rotate within its surrounding matrix, which remains stationary (Fig. 1B) [27]. The rotational motion changes the mechanical properties of the BM in a way that allows it to channel tissue growth along the anterior-posterior (AP) axis and create the elongated shape of the egg [29, 30]. Rotation always occurs around the AP axis, but it can be either clockwise or counterclockwise for a given egg chamber. Once a direction is chosen, however, the egg chamber rotates persistently in that direction for roughly two days until follicle cell migration stops at stage 8.

Egg chamber rotation is powered, in part, by lamellipodial protrusions that extend from the basal surface of each follicle cell [28]. These protrusions depend on the formation of a local branched actin network, under the control of the WASP family verprolin homolog regulatory complex (WRC), and their tissue-level alignment requires the atypical cadherin Fat2 (Fig. 1C) [31–33]. Fat2 localizes to the trailing edge of each follicle cell [34], where it acts *in trans* to create a stable domain of protrusive activity at the leading edge of the cell behind, and thereby orients all the cells’ protrusions in a common direction [32, 33]. Without Fat2, protrusions are unstable and unpolarized and rotation does not initiate [33].

Although we have learned a great deal about how Fat2 orients protrusions and other cellular features at the basal surfaces of the follicle cells when rotation is occurring at steady state [31–37], we still know very little about the symmetry-breaking mechanism that allows Fat2 to polarize the tissue for the first time and initiate rotation. Indeed, it has been notoriously difficult to capture the onset of rotation with live imaging, and the exact stage at which rotation begins is debated in the field [28, 37]. The mechanisms that ensure rotation always occurs around the egg chamber’s AP axis are similarly unknown.

In this paper we combine experiments with mathematical modeling to explore how Fat2 mediates chiral symmetry breaking in the follicular epithelium, and how the rotational axis is specified. By employing methods for long-term live imaging and for delaying the onset of rotation, we reveal the dynamic behaviors of the follicle cells as rotation initiates and show that symmetry breaking is a self-organized process. We then use our experimental observations as the basis for a theoretical model and find that a Fat2-based mechanochemical feedback loop and rigid-body dynamics of the egg chamber are sufficient to break chiral symmetry in the follicular epithelium and generate sustained rotation. Finally, we propose that mechanical constraints imposed by cell-cell interactions and tissue geometry define the rotational axis and ensure its stability over time. These findings provide a general framework for understanding how epithelial cells self-organize for rotational migration in the context of a complex, developing tissue.

## Results

### Egg chamber rotation begins within the germarium

To understand how chiral symmetry is broken in the follicular epithelium, it is essential to know when rotation begins under wild-type conditions. Although it is well accepted that rotation initiates shortly after an egg chamber forms, whether it occurs at stage 1, when the egg chamber resides in the germarium and is attached to the precursors of the stalk cells (pre-stalk), or stage 2 when it is separated from the germarium and fully encased in its own BM (Fig. 1D), has been difficult to discern using standard live imaging approaches. To overcome this barrier, we gently embedded young egg chambers in a fibrinogen-thrombin clot [38], which allowed us to perform continuous live imaging *ex vivo* for up to nine hours, and we visualized the BM using an endogenous GFP tag on the *α*2 chain of collagen IV (called Viking in *Drosophila*, Col IV-GFP) [39]. In all 12 samples examined, rotation was already occurring at an average rate of 0.2 µm/min at stage 1, before a BM had formed around the anterior of the egg chamber, and rotation persisted as the egg chamber transitioned to stage 2 (Fig. 1E-H, S1A,B; Movie 1). In 2 of the 12 samples, we even captured the onset of rotation and found that it reaches a constant rate within minutes after motion is first detected, which is then maintained for many hours (Fig. 1I, S1C; Movie 2). Altogether, these data show that the rotational migration of the follicle cells begins when the egg chamber is still in the germarium and that its onset has switch-like dynamics.

### Fat2 can mediate symmetry-breaking at multiple developmental stages

We next asked when Fat2 is competent to mediate symmetry-breaking in the follicular epithelium. Fat2’s symmetry-breaking function could be restricted to stage 1 when rotation normally begins, or Fat2 might be capable of mediating symmetry-breaking at multiple developmental stages. Prior work suggested that the former may be true [37]; however, this assertion is at odds with subsequent observations that Fat2 is continuously required to orient cellular protrusions throughout the rotational period [36], and that rotation can initiate at later stages in some mutant conditions [40]. To revisit this question, we delayed the onset of Fat2 expression and asked if it can mediate symmetry-breaking in older egg chambers.

We used two Gal4 drivers to deplete Fat2 with RNAi, *traffic jam-Gal4* (*tj-Gal4*) and *109-30-Gal4*, and assessed the level of depletion using an endogenous 3xeGFP tag on Fat2 (Fat2-3xGFP) [32]. *tj-Gal4* is expressed in the follicle cells throughout oogenesis [41] (Fig. S1D). Using *tj-Gal4* to express *fat2-RNAi* reduces Fat2 to undetectable levels during the entire migratory period and fully blocks rotation (Fig. 2A-D). By contrast, *109-30-Gal4* is primarily expressed in the follicle cells of the germarium and early egg chambers [42] (Fig. S1D). When *109-30-Gal4* is used to express *fat2-RNAi*, Fat2 is undetectable through stage 3, begins to be expressed in patches of cells at stage 4 (Fig. S1E), and returns to wild-type levels by stage 7 (Fig. 2A,B). Importantly, the pattern of rotational migration shows a similar temporal progression to that of Fat2 expression, with rotation blocked through stage 3, occurring slowly in some egg chambers at stages 4-5, and reaching normal speed by stage 7 (Fig. 2C,D; Movie 3). Thus, Fat2 can mediate the symmetry-breaking event needed to initiate rotational migration at various developmental stages.

**Figure 2.**
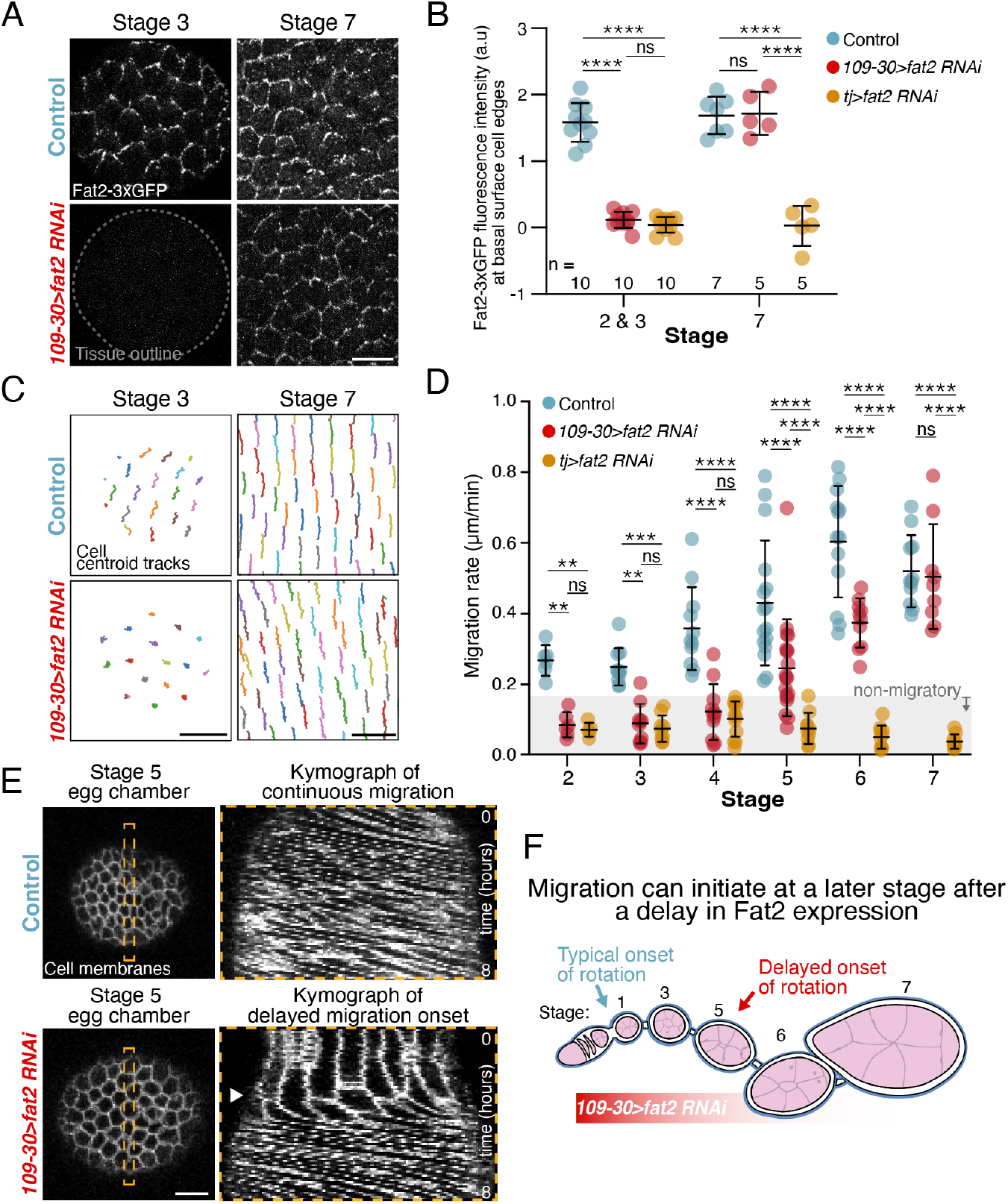
Fat2 can mediate symmetry-breaking at multiple developmental stages. **A)** Representative images of Fat2-3xGFP at the basal surface of the follicular epithelium. **B)** Quantification of Fat2-3xGFP intensity at basal surface cell edges. *109-30>fat2-RNAi* depletes Fat2 from the follicular epithelium at stages 2 and 3, but Fat2 levels are equivalent to controls by stage 7. Each data point represents one egg chamber. **C)** Representative centroid tracks of follicle cell movement over 10 minutes. **D)** Quantification of follicle cell migration rates over developmental time. *109-30>fat2-RNAi* blocks migration at early stages but migration is equivalent to controls by stage 7. Each data point represents one egg chamber. In order on graph, *n* = 6, 6, 5, 10, 10, 9, 11, 11, 12, 16, 19, 10, 11, 9, 9. **E)** Movie stills of stage 5 egg chambers with corresponding kymographs that show either constant migration or delayed onset of migration (arrowhead). **F)** Illustration summarizing how *109-30>fat2-RNAi* delays the onset of migration. For panels B and D, Two-way ANOVA with Tukey’s multiple comparisons test; ns, *p >* 0.05, \*\**p <* 0.01, \*\*\**p <* 0.001, \*\*\*\**p <* 0.0001. Bars represent mean *±* SD. Scale bars, 10µm.

We next asked if early depletion of other proteins required for symmetry breaking can also delay the onset of rotation. The WRC mediates the formation of cellular protrusions, and is required both to initiate and maintain follicle cell migration [28, 33]. We used *109-30-Gal4* to deplete the WRC component Abelson interacting protein (Abi) by RNAi and found that the onset of migration is delayed to the same extent as when Fat2 is depleted (Fig. S1F). These data show that the ability to initiate migration at a later stage is not exclusive to Fat2 and that it can likely be achieved by delaying the expression of any component of the symmetry-breaking machinery.

Finally, we asked if rotation initiation in these older egg chambers shows the same switch-like behavior seen at stage 1. We embedded ovarioles expressing *109-30>fat2-RNAi* in fibrinogen-thrombin clots for long-term live imaging and captured the initiation of rotation in multiple egg chambers that ranged from late stage 4 to early stage 6. Like the natural onset of rotation, this motion quickly reaches a constant rate, which is then maintained, showing that initiation is a rapid process regardless of when it occurs (Fig. 2E; Movie 4).

Altogether, these data show that there is no developmental pre-pattern governing when rotation initiates. Instead, it can occur at multiple stages, which suggests that the symmetry-breaking mechanism that polarizes the epithelium for migration is a self-organized process (Fig. 2F).

### Fat2 becomes planar polarized concurrent with the onset of rotation

The ability to delay the onset of rotation highlights the robustness of the symmetry-breaking process. It also provides a powerful method to probe the cellular dynamics that drive the transition into rotation. Because the number of follicle cells increases as the egg chamber develops, we can track the behaviors of more cells in a given experiment. With this new method, we first probed the dynamics of Fat2’s planar polarization to the trailing edge of each follicle cell, as it is the localization of Fat2 that orients all the cells’ protrusions in a common direction [33]. We previously proposed that the motion of the tissue itself localizes Fat2, suggesting that this is a mechanosesitive process [32]. However, this proposal came from experiments in which the epithelium was either rotating at steady state or rotation was fully blocked (Fig. S2A). Delaying the onset of migration now lets us ask if Fat2 becomes localized as rotation begins.

We used either *tj-Gal4* or *109-30-Gal4* to express *Abi-RNAi* and assayed the planar polarization of Fat2-3xGFP by determining the ratio of fluorescence intensity along leading-trailing cell edges versus lateral cell edges (Fig. 3A-C). When the epithelium is migrating, Fat2-3xGFP is enriched at leading-trailing edges at all stages examined. Conversely, blocking migration using *tj>Abi-RNAi* causes Fat2-3xGFP to localize uniformly around all cell edges [32]. When migration was instead delayed using *109-30>Abi-RNAi*, we saw a dynamic temporal pattern of Fat2 localization, with Fat2-3xGFP uniformly localized around cell edges at stages 3-4, weakly polarized to leading-trailing edges at stage 5, and robustly polarized by stage 6 (Fig. 3A-C). Hence, Fat2’s tissue-level polarization is undetectable-to-weak during the time when migration normally initiates in these epithelia and robustly polarized only after migration is well underway. The fibrinogen-thrombin mounting method that allows us to capture the migration initiation does not yet allow high-resolution imaging of the basal surface, so we cannot visualize Fat2 polarization in living tissue. Nonetheless, these data suggest that Fat2 functions within a mechanochemical feedback loop to promote rotation: Fat2 acts at the trailing edge of each cell to align the cells’ protrusions, and the resulting epithelial migration, in turn, localizes Fat2 to each cell’s trailing edge.

**Figure 3.**
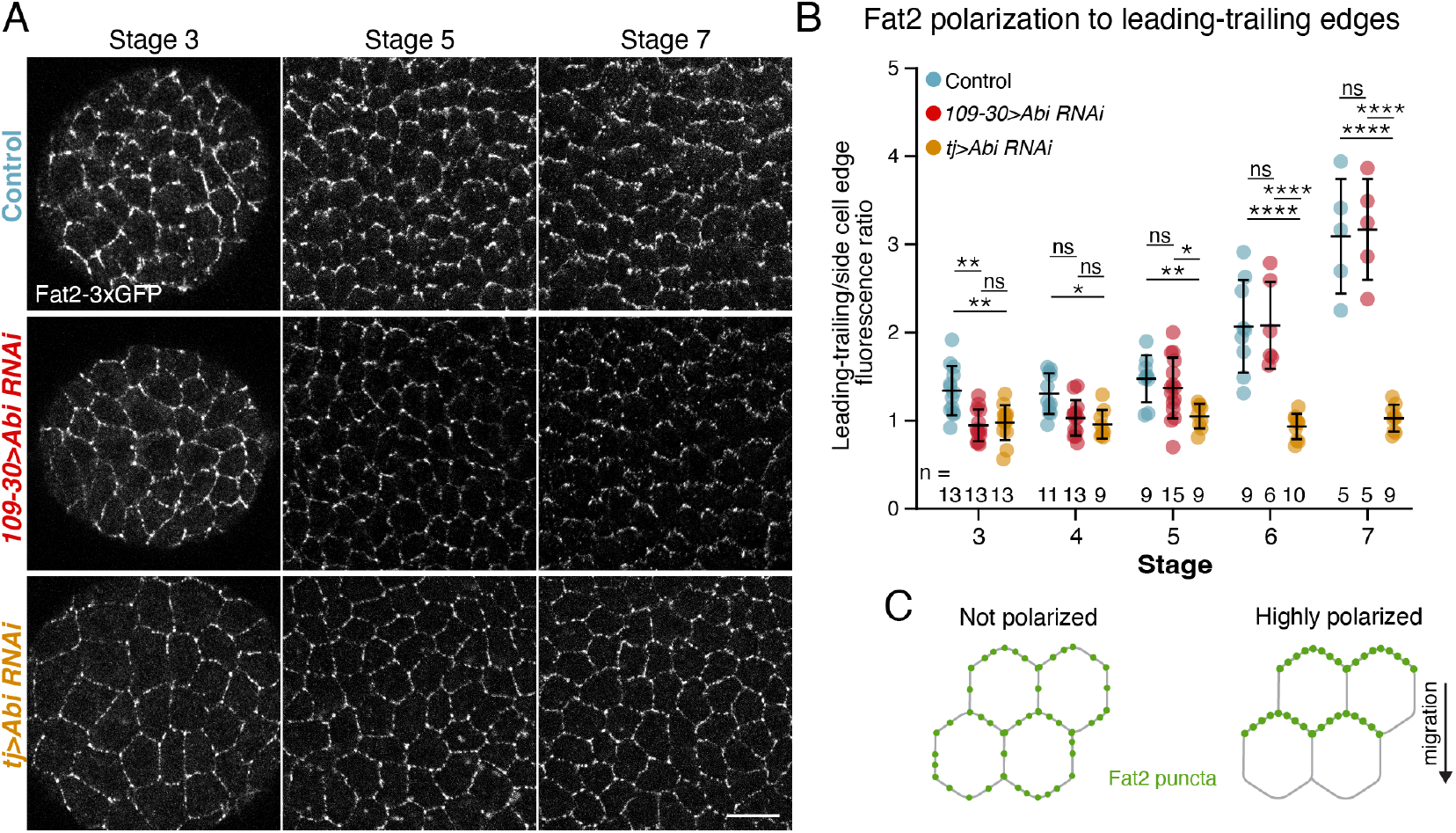
Fat2 becomes planar polarized concurrent with the onset of rotation. **A)** Representative images of Fat2-3xGFP at the basal surface of the follicular epithelium. **B)** Quantification of Fat2 polarization to leading-trailing cell-cell interfaces. Each data point represents the ratio of Fat2-3xGFP brightness at leading-trailing (horizontal) versus side (vertical) cell edges in one epithelium. Fat2 goes from unpolarized to polarized in *109-30>Abi-RNAi* with roughly the same timing as the onset of rotation in this background. Two-way ANOVA with Tukey’s multiple comparisons test; ns, *p >* 0.05, \**p <* 0.05, \*\**p <* 0.01, \*\*\*\**p <* 0.0001. Bars represent mean *±* SD. **C)** Illustration of non-migrating cells with uniformly distributed Fat2, versus migrating follicle cells with polarized Fat2. Scale bar, 10µm.

### Fat2 promotes local motility at the basal surface before rotation begins

We next used our delayed migration assay to visualize the dynamic movements of the follicle cells just before rotation begins and to determine how Fat2 influences these movements. For these experiments, we focused on stage 5, which is a *∼* 5-hr period [43] during which some *109-30>fat2-RNAi* epithelia are rotating, and some have not yet initiated rotation (pre-rotation) (Fig. 2D). For each epithelium, we took one 30-minute movie near the apical surface and one at the basal surface (Fig. 4C), and tracked the positions of cell centroids at both locations over time (Movie 5,Movie 6). In rotating *109-30>fat2-RNAi* epithelia, the apical and basal centroids of each cell move processively and in concert with one another, as expected. By contrast, in the pre-rotation *109-30>fat2-RNAi* epithelia, the apical cell centroids are largely static, while the basal centroids are highly dynamic (Fig. 4A,B; S2B,C; Movie 6). This local movement of the basal surface appears to be generated by the same protrusive activity that will eventually drive the rotational migration, with individual protrusions acting both cell-autonomously and non-autonomously to move the centroids of other cells (Movie 5). Notably, epithelia bearing the null mutation *fat2* ^*N103-2*^, which never migrate, show little centroid movement at their basal surfaces (Fig. 4A, S2B). Altogether, these data suggest that local motility at the basal epithelial surface may be a key feature of how rotation initiates and that Fat2 is required for this motility.

**Figure 4.**
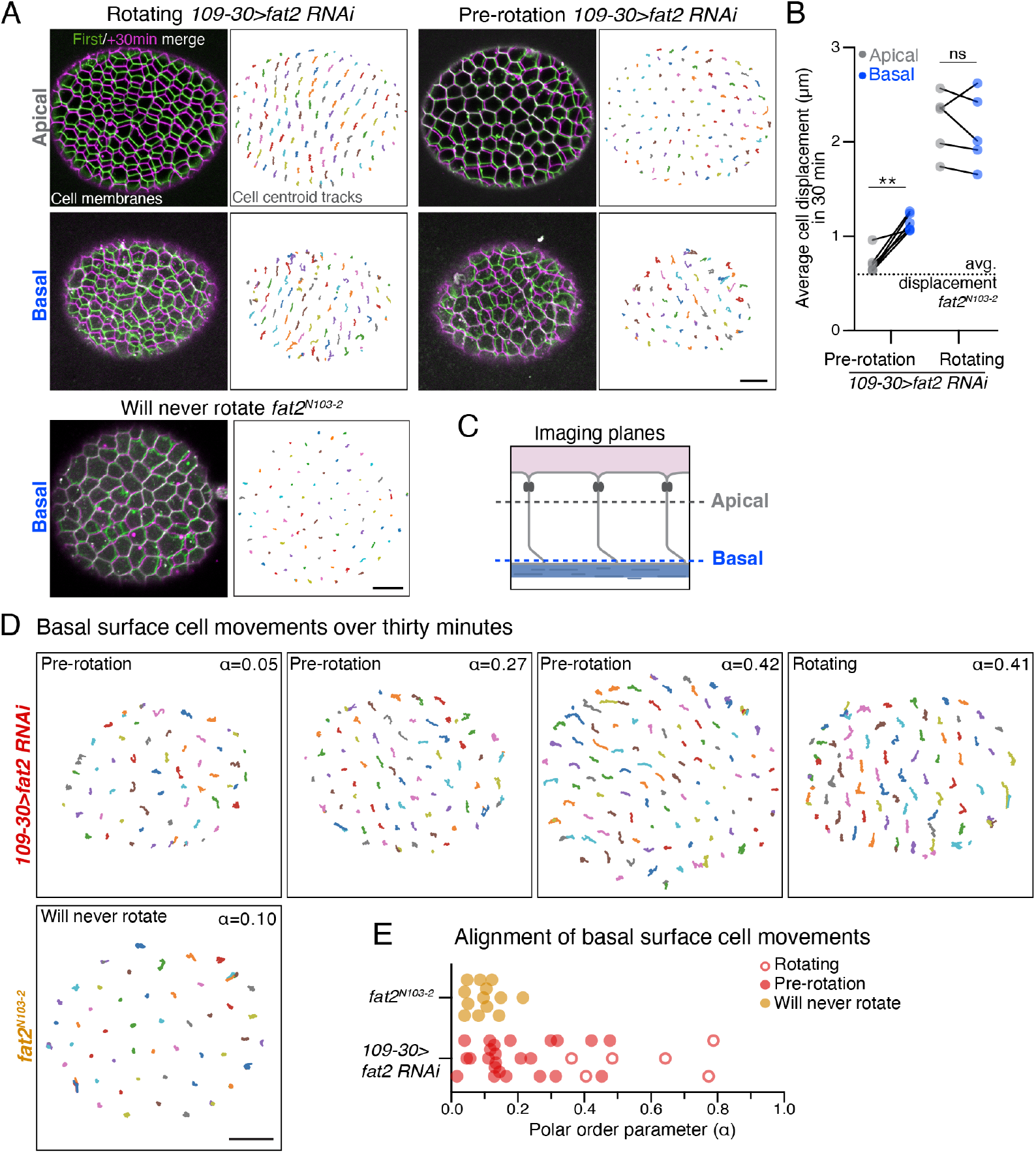
Fat2 promotes local motility at the basal surface before rotation begins. **A)** Representative images of a *109-30>fat2-RNAi* epithelium that is rotating, a *109-30>fat2-RNAi* epithelium pre-rotation, and a *fat2*^*N103-2*^ epithelium that will never rotate. Panels on the left show the overlay of the first (green) and last (magenta) frame of a 30-min movie (Movie 6). Panels on the right show cell centroid tracks from the same movie. **B)** Quantification of cell centroid displacement at apical versus basal surfaces in *109-30>fat2-RNAi* epithelia pre-rotation (*n* = 6) or that are already rotating (*n* = 5). Local motility at the basal surface precedes the onset of rotation. Each data point represents the average of cell displacements in one epithelium. Solid lines connect measurements from the same egg chamber. Dotted line at represents the average cell displacements from *fat2*^*N103-2*^ samples (Fig. S2). Paired t-test; ns, *p >* 0.05, \*\**p <* 0.01. **C)** Diagram denoting apical and basal imaging planes. **D)** Representative cell centroid tracks taken at the basal surfaces of five epithelia. The polar order parameter increases from left to right. **E)** Quantification of the polar order parameter for basal surfaces tracks of *109-30>fat2-RNAi* epithelia that are either rotating (red, open circles; *n* = 6) or pre-rotation (red, closed circles; *n* = 24), compared to *fat2*^*N103-2*^ epithelia (yellow circles; *n* = 13). Some pre-rotation epithelia have polar order parameters that exceed those of rotating epithelia, showing that the follicle cells can align their basal surface movements before rotation begins. Scale bars, 10µm.

We next investigated whether the local movements of the follicle cells’ basal surfaces become aligned before rotation begins by computing a polar order parameter *α* for each tissue (see methods). A polar order parameter of *α*=1 corresponds to all cell movements being perfectly aligned, while *α* = 0 corresponds to cell movements that are isotropic. Pre-rotation *109-30>fat2-RNAi* epithelia showed a range of tissue-level order, with some epithelia being largely isotropic, like *fat2* ^*N103-2*^ epithelia, and others showing varying degrees of alignment (Fig. 4D,E). Notably, in the *109-30>fat2-RNAi* condition, the extent of cell alignment in three pre-rotation epithelia exceeded that of some epithelia that were already rotating (Fig. 4E). We conclude that the follicle cells can align their basal surface movements before rotation begins and further speculate that each *109-30>fat2-RNAi* epithelium we analyzed represents a snapshot of the developmental trajectory a given epithelium takes from a disordered to an ordered state as it transitions into rotation.

It has long been known that the apical surfaces of the follicle cells adhere to the germ cells at the center of the egg chamber. This adhesion causes the germ cells to rotate in concert with the follicle cells [27] and likely limits neighbor exchange between the follicle cells over short time scales. In this way, the egg chamber rotates as a rigid body in which the primary motive forces are restricted to the basal epithelial surface where Fat2 is active. Our finding that Fat2-dependent cell motility is restricted to the basal surface before rotation begins suggests that this is also true during the symmetry-breaking process, which provides key insight into the mechanical state of the egg chamber when rotation initiates.

### Rigid-body dynamics and the mechanosensitive behavior of Fat2 can initiate rotation

The above results show that the follicle cells can start migrating from an unpolarized state and that rotation initiation likely occurs with rigid-body dynamics. We further hypothesize that Fat2 operates within a mechanochemical feedback loop to promote rotation, in which Fat2 acts at the trailing edge of each cell to align the cells’ protrusions, and the resulting tissue motion localizes Fat2 to the cells’ trailing edges. Here, we develop a minimal theoretical model to elucidate whether these conditions can recapitulate rotation initiation.

We modeled the egg chamber as an ellipsoidal rigid body (Fig. 5A). Crawling forces are generated by lamellipodial protrusions at the basal epithelial surface, which we model as point forces **b**_*i*_ generated by each follicle cell *i ∈* 1, 2,..*N* at positions **r**_*i*_ (Fig. 5B). Throughout, we denote vectors and matrices in bold and scalars in regular font. The BM provides the substrate for cell crawling but also generates resistance to motion. We model this resistance as a viscous drag force with viscous coefficients *µ*_*b*_ for motion tangential to the follicle cell-BM interface (Fig. 5B). The BM elastically resists motion perpendicular to the egg chamber-BM interface. Intuitively, the egg chamber can be thought of as a rigid ellipsoid elastically confined by a surrounding BM. Any motion distinct from rotation around the long axis generates restoring elastic forces/torques. Given estimates of the stiffness of the BM (*≈* 50 kPa) [30, 44] and traction stress generated by other crawling cells *in vivo* (order-of-magnitude *≈* 10Pa [45]), we infer that this elastic response strongly constrains translational motion. Furthermore, for an ellipsoidal egg chamber (eccentricity *e >* 1), elastic confinement restricts rotation to the long axis, which reflects what we observe with the delayed migration assay (See Sec. S1.1.1 for details). These observations allow us to reduce the full equation of motion for the egg chamber to a single equation describing rotation around its long axis. In the overdamped limit (negligible inertia), the non-dimensional equation for the rotation of the egg chamber along the long axis (Sec. S1.1) is

**Figure 5.**
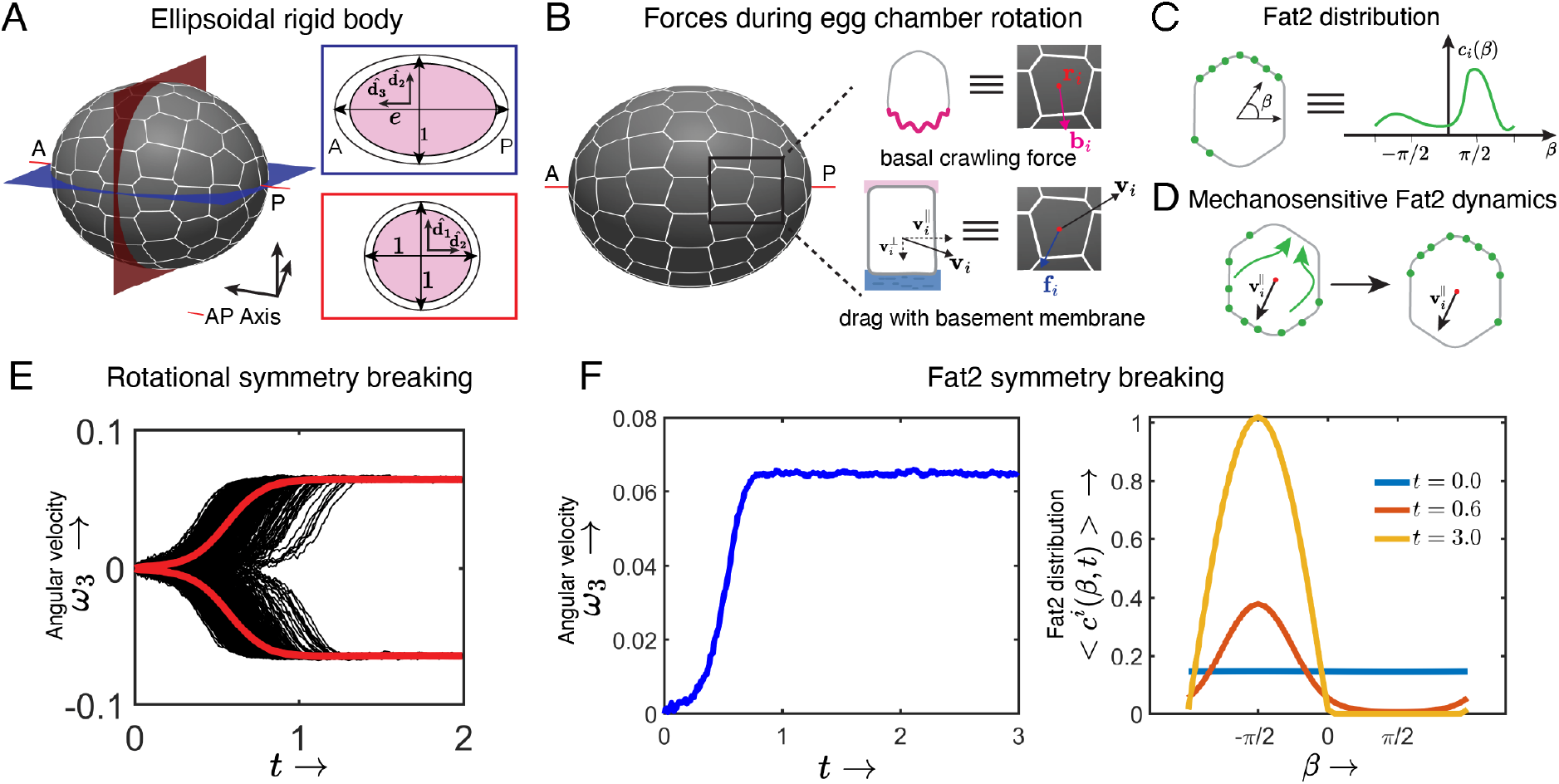
Rigid-body dynamics and the mechanosensitive behavior of Fat2 can initiate rotation. **A)** To model the delayed migration assay, the egg chamber is represented as an ellipsoidal rigid body with eccentricity *e*. The dimensions of the egg chamber are rescaled with the radius of the equatorial cross section. **B)** The crawling force *b*_*i*_ models protrusive activity at the basal epithelial surface and the viscous drag *f*_*i*_ models the interaction between the epithelium and the BM for cell *i*. **C)** The Fat2 distribution is modeled by *c*_*i*_(*β, t*) where *β* is the angular direction in the tangent space. **D)** Fat2 dynamics are mechanosensitive, whereby Fat2 is recruited to each cell’s trailing edge with respect to the migration direction *v*_*i*_. **E)** The model breaks rotational symmetry, generating either clockwise (*ω*_3_ *<* 0) or counter-clockwise (*ω*_3_ *>* 0) rotation about the AP axis. *N* = 1000 model trajectories starting from an isotropic initial condition (*c*_*i*_(*β*, 0) = 1*/*2*π, b*_*i*_ = 0). **F)** Representative model trajectory with symmetry broken, resulting in sustained counterclockwise rotations and the corresponding evolution of the cell-averaged Fat2 distribution *⟨c*_*i*_(*β, t*)*⟩*. The Fat2 distribution evolves from an isotropic configuration to a polar configuration centered around *β* = *−π/*2. The time evolution of the quantities in F is given in Movie 7.

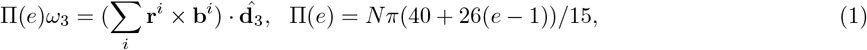

where the angular velocity vector is 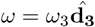 and 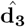 is the unit vector pointing to the anterior pole along the AP axis (Fig. 5A).

Next, we modeled the interplay between the planar polarization of Fat2 and protrusions. We account for the Fat2 distribution as a concentration field *c*_*i*_(*β, t*), where *β* is the angular coordinate in the tangent plane at **r**_*i*_ (Fig. 5C). Although Fat2 acts *in trans* to orient protrusions, its function is restricted to the cell-cell interface where it resides [32]. Given this local effect, we simplify the model by representing Fat2’s function as being cell-autonomous. A minimal model of Fat2’s effect on follicle cell protrusions is

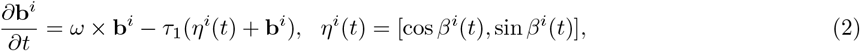

where *τ*_1_ = *t*_*c*_*/γ* is the ratio of the characteristic time-scale of rotation initiation to the time-scale over which the crawling force changes directions in response to a change in Fat2 distribution. The effect of the Fat2 distribution *c*_*i*_(*β, t*) on crawling force dynamics is encoded through *η*_*i*_(*t*) on the tangent plane at **r**_*i*_, where *β*_*i*_(*t*) is a stochastic variable sampled from the distribution proportional to *c*_*i*_(*β, t*) (Sec. S1.2). Eq. (2) models the alignment of the protrusions at the cell edges opposite to those with Fat2 enrichment (Fig. 5C,D).

Last, we modeled the mechanosensitive dynamics of Fat2, in which Fat2 localizes to cell edges opposite to the local cell migration direction **v**_*i*_ (Fig. 5D and Sec. S1.2.1)

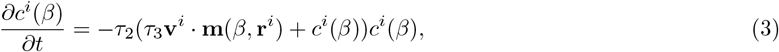

where *τ*_2_ = *ϵ*_0_*t*_*c*_*c*_*c*_, *τ*_3_ = *ϵ*_1_*a/t*_*c*_*c*_*c*_, *c*_*c*_ is the characteristic concentration scale of Fat2, *ϵ*_0_ is a characteristic rate of Fat2 dynamics, *ϵ*_1_ sets the relative strength of the mechanosensitivity compared to saturation dynamics, *a* is the radius of the equatorial cross-section of the egg chamber and **m**(*β*, **r**_*i*_) is a unit tangent vector along the angular direction *β* at **r**_**i**_ (Sec. S1.2.1).

Consistent with observations of pre-rotation egg chambers in the delayed migration assay, we solve Eqns (1-3) with initially uniform (i.e. symmetric) Fat2 (*c*_*i*_(*β, t*_0_) = 1*/*2*π*) and crawling force distribution (**b**_*i*_(*t*_0_) = 0) (See Sec. S1.5 for details). For parameter selection (ie, *τ*_1*−*3_) and sensitivity analysis see Sec. S1.3. Using these initial conditions and parameters, the model breaks symmetry and generates sustained rotations (Movie 7). These rotations occur about the AP axis, with equal probability of clockwise (*ω*_3_ *<* 0) and counterclockwise (*ω*_3_ *>* 0) directions, consistent with experiments (Fig. 5E). The Fat2 distribution and crawling force transition from an isotropic to a polar distribution centered around *β* = *π/*2 for clockwise and *β* = *−π/*2 for counter-clockwise rotations. These observations reflect Fat2 being localized to the cells’ trailing edges and the progressive alignment of cell protrusive activity as rotation initiates. Fig. 5F shows one simulation iteration that undergoes counterclockwise (*ω*_3_ *>* 0) rotational symmetry breaking and the cell-averaged Fat2 distribution *< c*_*i*_(*β, t*) *>*, transitioning from an isotropic to a polar distribution centered around *β* = *−π/*2. Our results remain robust to large parameter changes (Sec. S1.3).

Here, egg chamber geometry sets the rotation axis and the rigid body dynamics provides the global synchronizing mechanical cue for follicle cells, while the mechanosensitive behavior of Fat2 transduces this cue to align cell polarity. The transduction of the global rotational cue into the Fat2 distribution further promotes crawling force in the direction of rotation, thereby forming positive feedback between rotation, Fat2 distribution, and cell crawling force (Fig. 7).

### Mechanical constraints specify the rotational axis

Wild-type egg chambers always rotate around the AP axis [27]. Notably, this is also true when the onset of rotation is delayed using *109-30>fat2-RNAi*. In the above model, elongation of the egg chamber along its AP axis (*e >* 1) (i.e., egg chamber geometry), combined with elastic confinement from the BM, ensures that rotation occurs around the AP axis. However, egg chambers are thought to be spherical during the earliest developmental stages when rotation normally begins. Simulating the above model with a sphere (*e* = 1), the epithelium still breaks symmetry, but rotation is not restricted to any specific axis (See Sec. S1.4). We therefore asked how the rotational axis is specified under wild-type conditions.

Having shown that rotation begins at stage 1, we looked more closely at the shape of the egg chamber within the germarium. Although stage 1 egg chambers are mostly round, the follicle cells at the anterior pole form a flattened interface with the pre-stalk cells that will eventually separate individual egg chambers after they bud from the germarium (Fig. 6A, S2D) [46]. Live imaging of rotating stage 1 egg chambers showed that these anterior-most follicle cells rotate with the rest the tissue and slide past the pre-stalk cells, which remain stationary (Fig. 6B, S2E; Movie 8). This led us to hypothesize that a mechanical interaction between the egg chamber and the pre-stalk cells could constrain the direction of rotation.

**Figure 6.**
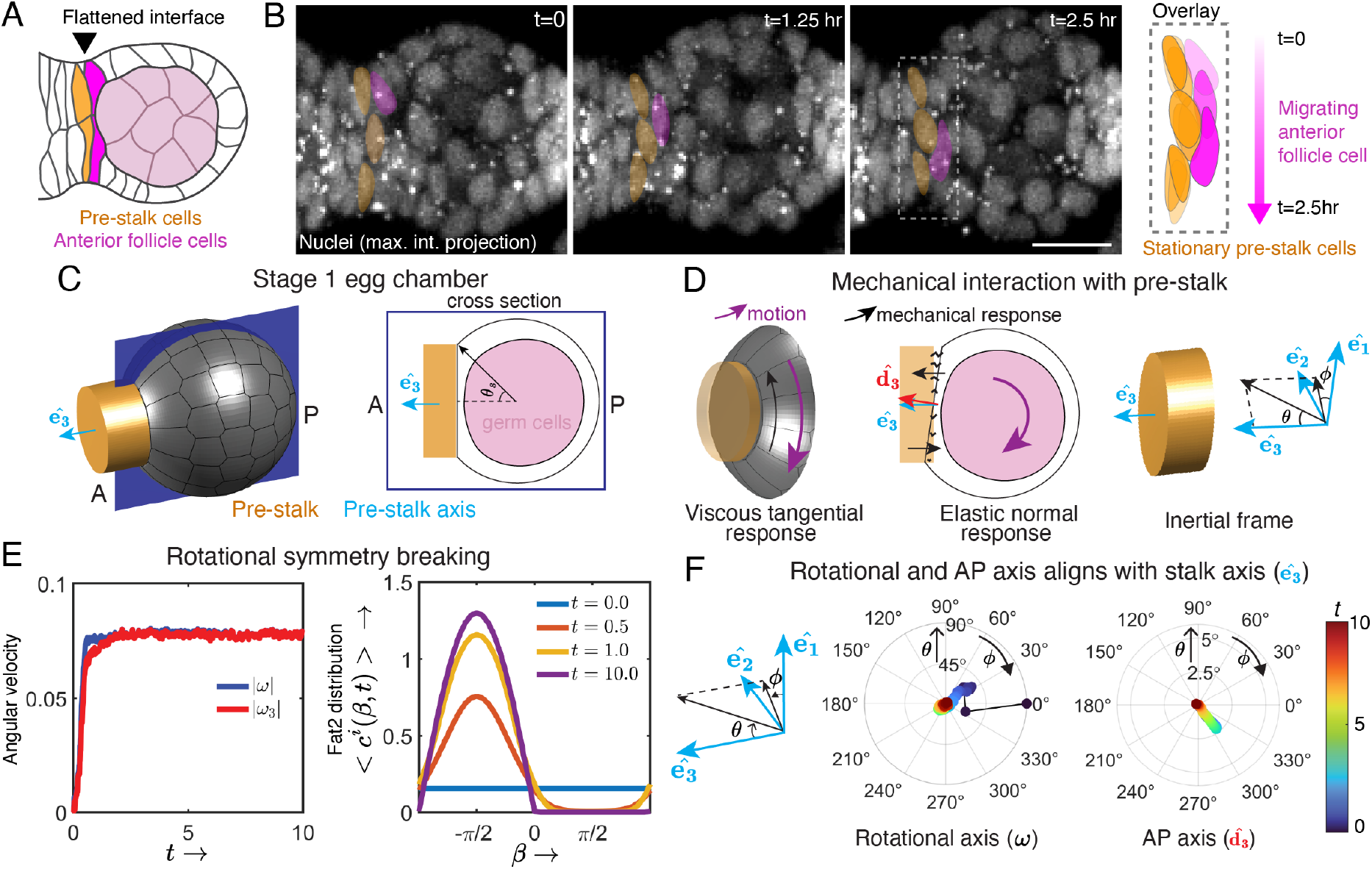
Mechanical constraints specify the rotational axis. **A)** Illustration of a transverse section through a rotating egg chamber at stage 1, highlighting the shape of the interface between a subset of pre-stalk cells (gold) and the anterior-most follicle cells (magenta) in the germarium. **B)** Movie stills of maximum intensity projections of nuclei generated from a 2.5-hour movie (Movie 8). Stationary pre-stalk cell nuclei are pseudocolored gold, migrating anterior follicle cell nuclei are pseudocolored magenta. The overlay shows the change in the cells positions at these time points. **C)** The polar angle *θ*_*s*_ quantifies the extent of contact between the pre-stalk cells and the egg chamber. The unit vector 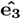 is along the pre-stalk axis, which aligns with the egg chamber’s AP axis. **D)** The motion of the egg chamber tangential (normal) to the pre-stalk/egg chamber interface generates viscous (elastic) resisting forces. Rotational motion of the egg chamber is described in the inertial (i.e. fixed) frame 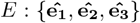. The orientation of a vector is represented in spherical coordinates (*θ, ϕ*). **E)** The rotational symmetry-breaking dynamics of a stage 1 egg chamber is described using 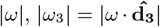 and average Fat2 distribution *⟨c*_*i*_(*β, t*)*⟩*. **F)** The evolution of the orientation of the rotational axis *ω* and the AP axis 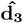 is described using the spherical coordinates (*θ, ϕ*) in the inertial frame *E*. The time evolution of the quantities (F-G) is given in Movie 9.

To test this hypothesis, we extended our model to account for the mechanical interaction with pre-stalk cells. The areal extent of the egg chamber/pre-stalk interface is quantified by the polar angle *θ*_*s*_ (Fig. 6C). The interaction is assumed to be elastic for motion normal to this interface. For motion tangential to the interface, instead, we assume a viscous drag force (Fig. 6D). We use the same Fat2 and protrusion dynamics as in Eq. 2 and Eq. 3. By numerically simulating this extended model (See Sec. S2 for more details), the egg chamber breaks symmetry, generating sustained rotations about the AP axis (Fig. 6E,F; Movie 9). There are three relevant axes here. The stationary pre-stalk axis (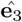) and two dynamic axes: the AP axis (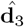) that comoves with the egg chamber, and the egg chamber rotational axis (*ω*) (Fig. 6D). The AP axis is defined by the axis normal to the flattened interface with the pre-stalk cells. The time taken to align the rotational axis with the pre-stalk axis depends on the strength of the elastic interaction with the pre-stalk cells. Stronger elastic interactions induce faster alignment.

Therefore, the mechanochemical feedback loop that breaks symmetry, combined with mechanical interaction with the pre-stalk cells, can generate sustained rotation about the AP axis in stage 1 egg chambers. In this natural condition, the geometry of the pre-stalk/egg chamber interface sets the rotational axis as rotation initiates. At later stages, we envision that the elongated shape of the egg chamber ensures that the rotational axis remains stable over time.

## Discussion

This work sheds light on two key aspects of egg chamber rotation in *Drosophila* - how rotation initiates and how the rotational axis is specified. We show that the follicle cells initiate rotational migration at stage 1, when they are still in the germarium, and that chiral symmetry breaking is a self-organized process. Our data and modeling further suggest that a Fat2-based mechanochemical feedback combined with the rigid-body motion of the egg chamber is sufficient to both initiate rotational migration and make it self-sustaining. Finally, our work suggests that rotation about the AP axis is specified by a tissue-geometry-mediated mechanical interaction between the anterior-most follicle cells and the pre-stalk cells at stage 1, and that the axis is then maintained by the progressive lengthening of the egg chamber over time (Fig. 7). The central features of our proposed mechanism for egg chamber rotation are detailed below.

**Figure 7.**
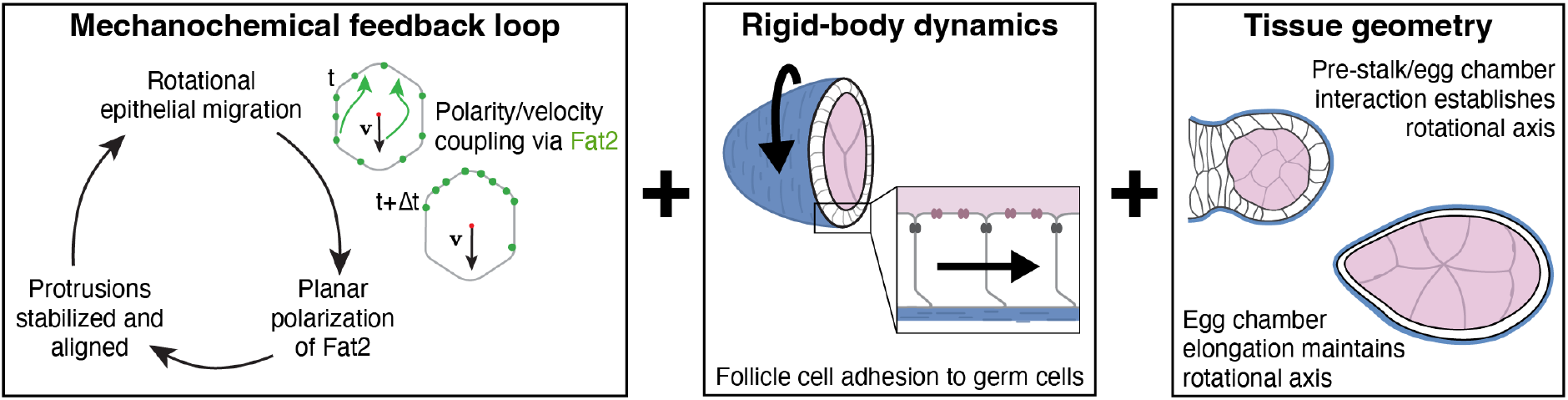
Proposed biophysical mechanism for egg chamber rotation. Fat2 operates in a mechanochemical feedback loop, in which cell motion polarizes Fat2 to cells’ trailing edges, which in turn stabilizes and aligns individual cell crawling forces. Rigid-body dynamics of the egg chamber synchronize the individual movements of the follicle cells for collective migration. Tissue geometry provides a mechanical cue that ensures rotation around the AP axis. Synergy between the mechanosensitive behavior of Fat2, egg chamber rigid-body dynamics, and egg chamber geometry initiates and maintains the rotational migration of the follicle cells.

We propose that Fat2 operates in a mechanochemical feedback loop to initiate and maintain the rotational migration of the follicle cells. By delaying the onset of Fat2 expression, we found that Fat2 can initiate rotation at multiple developmental stages and that initiation can occur *ex vivo*, showing that chiral symmetry breaking is a self-organized process. Moreover, because follicle cell number [47–49] and egg chamber volume [43] both increase roughly 10-fold over the stages at which we have seen rotation initiate, symmetry-breaking is robust to these factors. We also found that Fat2 becomes planar polarized concurrent with the onset of rotation. This observation, combined with prior data [32] and current modeling, suggests a positive feedback loop in which Fat2 aligns the crawling forces of individual cells and the resulting rotational motion polarizes Fat2 to the cells’ trailing edges. We envision that Fat2 aligns the crawling forces by acting *in trans* to create a stable domain of protrusive activity at the leading edge of the cell behind [32, 33]. However, Fat2 may also act on other components of the migration machinery [31, 34, 35, 37]. Elucidating how the resulting collective motion, in turn, leads to polarized localization of Fat2 is an important area for future research.

We further propose that the rigid-body dynamics of the egg chamber synchronizes individual follicle cell movements for collective migration. Unlike most epithelial cells, which have free apical surfaces, the apical surfaces of the follicle cells tightly adhere to the germ cells [27]. This adhesion rigidifies the tissue and limits neighbor exchange on short time scales. Consistent with this mechanical constraint, we found that cell motility is restricted to the basal epithelial surface before rotation begins and that this local motility depends on Fat2, likely through its effect on cell protrusive activity [33]. For rotation to initiate, the movements of the individual cells must become aligned in a global migration direction, which we also observed. Our model posits that rigid-body dynamics provides this global synchronizing cue. Indeed, combining the rigid-body assumption with the mechanosensitive behavior of Fat2 described above (polarity-velocity coupling) [11, 50] is sufficient to break symmetry and generate sustained egg chamber rotations. Exploiting rigid-body dynamics, our model yields closed-form parametric expressions that relate tissue geometry (e.g., eccentricity, pre-stalk interface size and shape) to mechanical outputs, offering insight into how geometry influences tissue dynamics. Overall, our biophysical mechanism extends prior models of symmetry breaking proposed for rotating epithelial spheroids, based on cell flocking behaviors through Viscek-like polarity alignment [17, 18] or cell substrate curvature [22]. Beyond the egg chamber, we anticipate that the same principles can drive chiral symmetry breaking even when the rigid-body assumption is relaxed to allow elastic deformations of the rotating tissue—a generalization we will pursue in future work.

Another major finding of this work is that tissue-geometry-mediated mechanical constraints may ensure that rotation occurs around the AP axis. It has been proposed that two pairs of specialized follicle cells (polar cells) define the rotational axis [51], as they are located at the anterior and posterior poles of the egg chamber. However, the polar cells have not yet differentiated when rotation initiates at stage 1 [52]. Instead, the anterior-most follicle cells directly contact the pre-stalk cells and can remodel their contacts along this flattened interface as rotation initiates. Our model suggests that an elastic interaction between these two cell types is sufficient to specify the rotational axis at this stage. Delaying the onset of migration revealed that rotation also occurs around the AP axis when it initiates at stages 4-6. Here, slight elongation of the egg chamber, combined with elastic confinement from the BM can specify the rotational axis (Sec. S1.1.1). At these stages, polar cells would be required because they control the early, rotation-independent phase of tissue elongation [53]. We speculate that progressive lengthening of the tissue maintains the rotational axis in wild-type egg chambers, ensuring its stability over roughly two days when this motion occurs *in vivo*.

Importantly, three features of our proposed mechanism for egg chamber rotation require interactions between the migrating follicle cells and other non-migratory cell types – the germ cells, the pre-stalk cells and the polar cells. This contrasts with many *in vitro* systems of rotational epithelial migration that focus on uniform monolayers of migrating cells [10, 14–17, 22]. One notable exception is human mammary organoids, which have rotating spherical acini attached to a central system of ducts [23]. Unlike isolated epithelial spheres that do not maintain a coherent rotational axis, these acini rotate persistently around the ducts in a manner that is remarkably like that of a stage 1 egg chamber rotating around the pre-stalk cells. Thus, interactions with non-migratory cell types may provide a general mechanism to constrain rotational epithelial migrations, even though rotational migration, itself, is a spontaneous and self-organized process.

## Methods

### Fly genetics, sources and care

*Drosophila melanogaster* were fed cornmeal molasses agar food and raised at 25°C. Experimental females were collected 0-3 days post-eclosion and then aged in the presence of males with yeast before dissection. In all RNAi experiments using *109-30-Gal4* or *tj-Gal4*, experimental and control females were kept at 29°C for 48 hours prior to dissection, otherwise experimental females were kept at 25°C. The genotypes used in each experiment are listed in Supplementary Table 2 and indexed by figure panel.

**Table 1:**
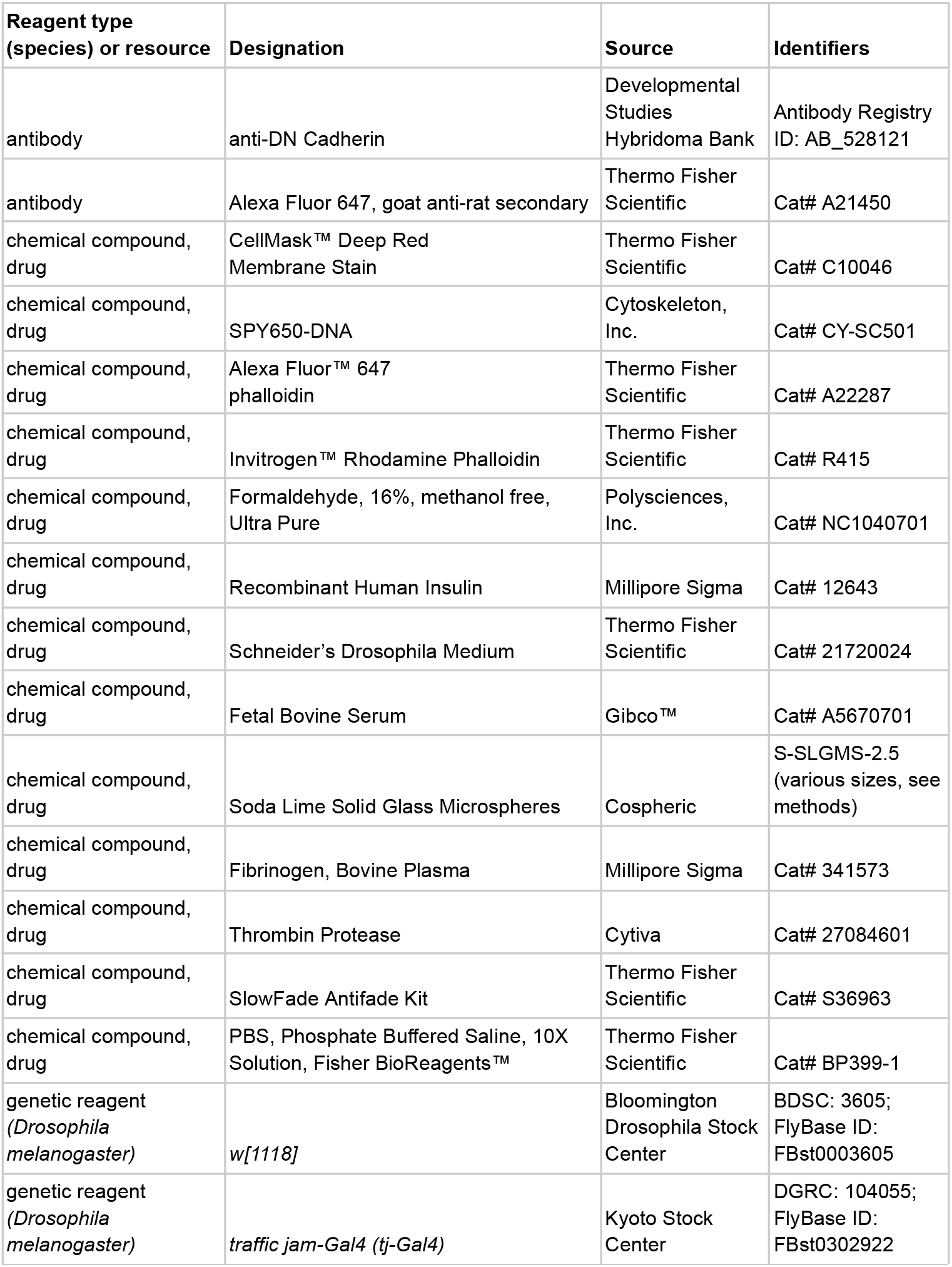
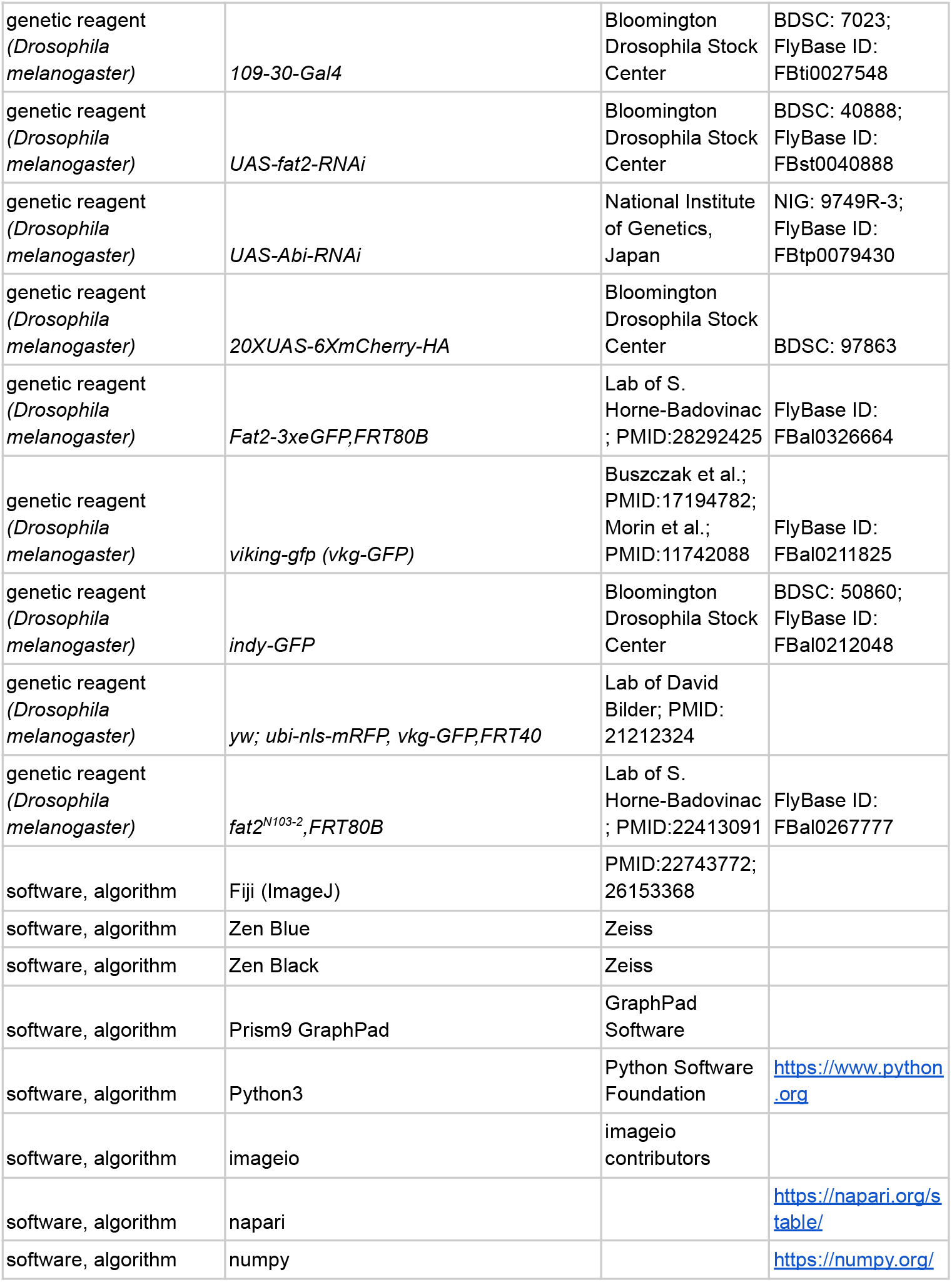
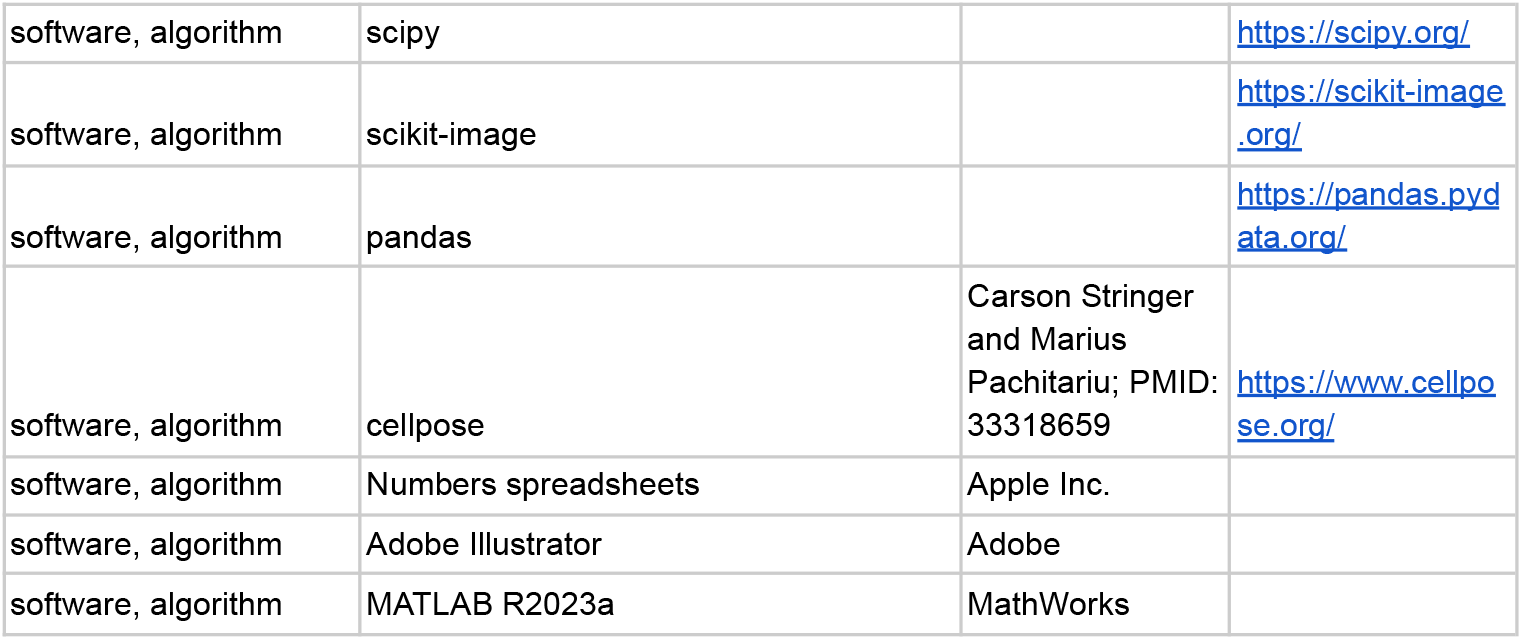
Resources and reagents.

**Table 2:**
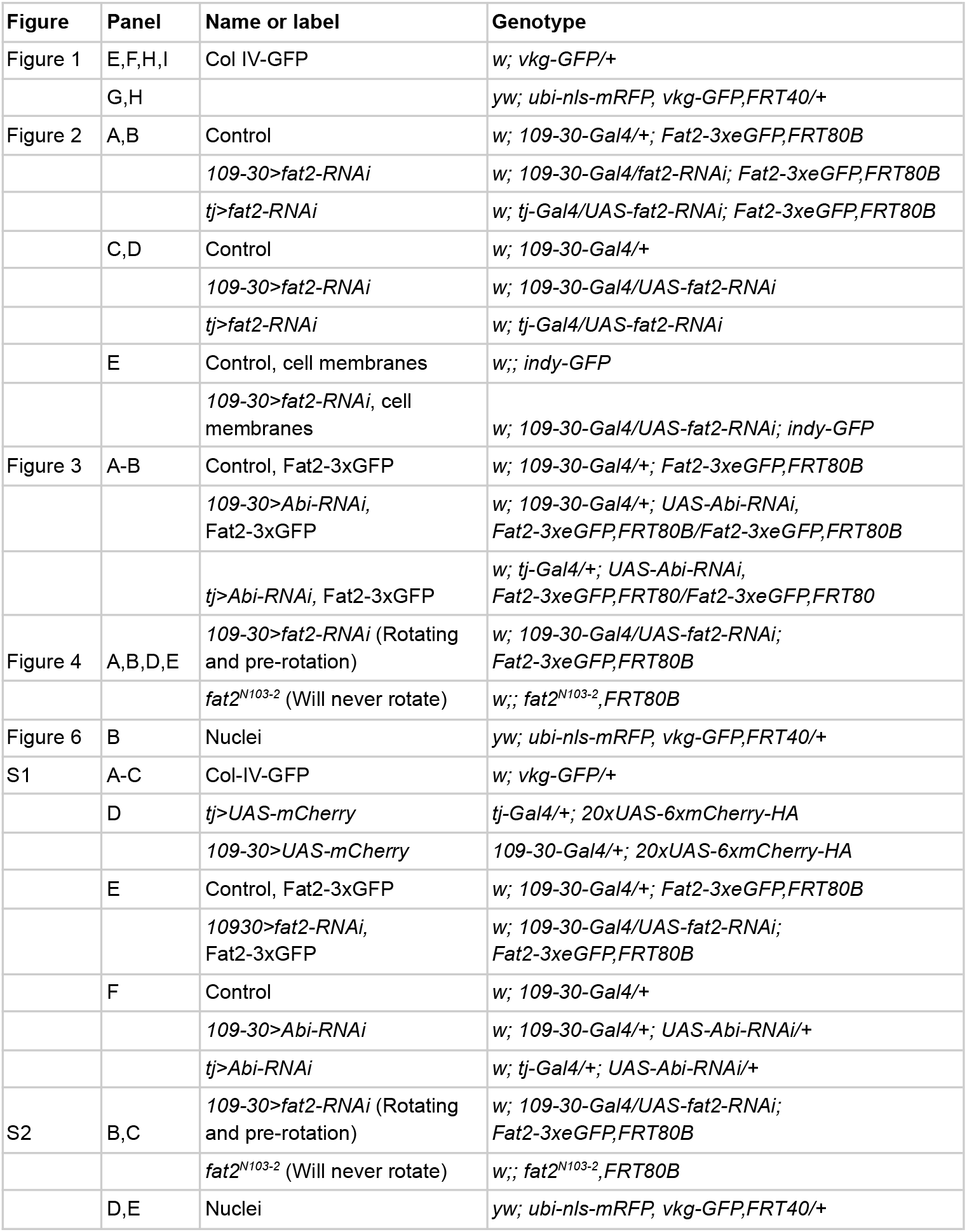
Experimental genotypes.

### Sample preparation

#### Egg chamber dissection

Ovaries were dissected into live imaging media (Schneider’s *Drosophila* medium containing 1X Penicillin-Streptomycin + 15% fetal bovine serum + 200µg/mL insulin, [54]) in one well of a glass spot plate using forceps. Ovarioles were carefully removed from the muscle sheath to not damage the stages of egg chambers being analyzed. For standard live imaging, any egg chambers older than the most mature egg chamber to be analyzed were trimmed from the ovariole and discarded. This was done by slicing through the stalk connecting the adjacent egg chambers with a 27-gauge needle. For extended live imaging, any egg chambers older than stage 9 were removed in the same manner. For a detailed protocol and movies of the dissection process see Cetera et al., 2016 [55].

#### Sample preparation for standard live imaging

Following dissection and trimming of the ovarioles, the tissue was transferred to another well in the spot plate containing live imaging media with CellMask (1:500) for 8 minutes and then washed with fresh live imaging media to remove excess dye. Stained ovarioles were transferred in 16µl of media to a slide, and glass spacer beads were added to the drop of media to limit egg chamber compression by the coverslip. The size of the beads varied based on the egg chamber stage of interest (51/53µm for stage 6-8, 40µm for stage 5-6, 30µm for stage 4-5, 25µm for stage 3-4, 15µm for germaria-stage 2). A coverslip was then placed on top of the ovarioles and beads and sealed with petroleum jelly to prevent evaporation. Imaging was performed with this setup for up to 1 hour.

#### Sample preparation for extended/overnight live imaging

Here, we adapted a method based on Wilcockson and Ashe, 2021 [38] to perform extended or overnight live imaging of egg chamber rotation. Following dissection and trimming of the ovarioles, the tissue was transferred to another well of the spot plate containing live imaging media with 1x SpyDNA for 15 minutes. During this 15-minute incubation, 200µl of prewarmed (37°C) live imaging media without insulin was added to an Eppendorf tube containing 0.002 g of fibrinogen (final concentration 10 mg/ml), vortexed briefly, then placed back at 37°C for the remainder of the staining period. Of note, increasing the volume of fibrinogen/media mixture or over-vortexing often resulted in premature clotting. After staining, the fibrinogen mixture was transferred to an adjacent well in the spot plate. Ovarioles were transferred to the fibrinogen mixture, pipetting slowly up and down to ensure complete incorporation, then transferred in 10µl to an imaging dish (MatTek, 35mm petri dish, 14mm microwell). Thrombin (10U/µl) was added to the drop of fibrinogen in two 0.5µl increments (1.0µl total), and left for 10 minutes. If the clot had not formed after 10 minutes another 0.5µl of thrombin was added. The clot was determined to be sufficiently formed if an eyelash could not break the surface tension of the clot-droplet, or polymerization was visible. In some, but not all samples, a Millicell culture insert (Sigma, 12 mm diameter, 8µm membrane pore size) was placed on top of the clot to prevent Z-drift but this was not a necessary step. The clot was then covered in 0.3 ml of live imaging media with insulin, a wet Kimwipe was added to the side of the dish to maintain humidity and the lid was placed on the dish. The z-stack acquisition feature was used to capture at least half the volume of egg chambers imaged. Tissues were imaged overnight for up to 13 hours; however, egg chamber rotation often slowed after 8-9 hours of imaging.

#### F-actin staining and immunostaining of fixed tissues

All fixed tissues were stained for F-actin (phalloidin) to facilitate the staging of egg chambers and N cadherin (anti-DN cadherin) to facilitate the segmentation of cells in the follicular epithelium for quantitative analyses. Dissected ovarioles were fixed in a 4% EM-grade formaldehyde in phosphate buffered saline (PBS) plus 0.1% Triton X-100 (PBT) for 15 minutes and then washed 3×5 minutes in PBT. Egg chambers were incubated with primary antibodies (anti-DN cadherin, 1:200) in PBT either overnight at 4°C or for 2 hours at room temperature while rocking, then washed 3×5 minutes in PBT. Ovarioles were then incubated in secondary antibodies (1:200) with the addition of either TRITC phalloidin (1:200) or Alexa Fluor 647 phalloidin (1:100) overnight at 4°C or 2 hours at room temperature while rocking. Ovarioles were then washed 3×5 minutes in PBT and mounted in 30µL SlowFade Diamond antifade on a slide using a 22×50 mm # 1.5 coverslip, sealed with nail polish and stored at 4°C until imaged.

### Microscopy

All live and fixed imaging was performed at room temperature using laser scanning confocal microscopy. Extended live imaging was performed using a Zeiss LSM 880 inverted microscope running Zen Black software with a 40x PlanAPOCHROMAT 1.3NA oil immersion objective (Fig. 1, 2E, 6A,B). All other images and movies were collected using a Zeiss LSM 800 upright microscope running Zen Blue software with a 63x Plan-APOCHROMAT 1.4NA oil immersion objective. All images were collected such that the egg chambers’ AP axes aligned with the horizontal (x)image axis, except in cases where this alignment occurred post-imaging in Fiji as described below.

### Analysis of live imaging data

#### Kymograph generation

To produce the kymographs shown in Fig. 1I, 2E, and S1C movies were aligned in Fiji (imageJ) so that the egg chambers’ AP axis corresponds with the horizontal (x) image axis. A maximum-intensity projection was generated to capture the height of follicular epithelium (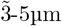) for the whole duration of the timelapse. In Fiji, the straight line tool (1 pixel) was used to draw a vertical line and the KymoResliceWide plugin was used to create the final kymograph.

#### Line-scans of Col IV-GFP

Timelapse movies of a central optical plane through stage 1 egg chambers were rotated in Fiji so that the AP axes corresponded with the horizontal (x) image axis. In Fiji, straight lines of 0.28µm (1 pixel) thickness were drawn vertically through the anterior-most pole of the egg chamber where the anterior BM will eventually form as determined from the movie. The average intensities of Col IV-GFP were measured at the beginning of the timelapse using the Plot Profile function in Fiji.

#### Cell segmentation and centroid tracking of live tissues

In brief, timelapse movies of CellMask stained epithelia were segmented in each frame using the pretrained ‘cytoplasm’ model in Cellpose [56] and tracked by linking regions of high overlap in consecutive frames. Napari tools were used to manually correct any cell segmentation and tracking errors. To account for XY drift, Python scikit-image and scipy libraries were used to generate a mask of the tissue. Tissue masks were centered in each frame based on their centroid position, then movies were aligned to centered tissue masks. Scikit-image regionprop was then used to determine the individual cell centroid positions at each frame of the movie. See Williams et al., 2022 [33] and Williams and Horne-Badovinac, 2023 [36] for more detailed protocols. See also ‘Code’ section below.

#### Migration rate data collection and analysis

For quantification of follicle cell migration rates as shown in Fig. 2D and S1F, ovarioles were stained with Orange or Deep Red CellMask (1:500) and mounted for standard live imaging. Timelapse movies were collected at a single Z plane just below the apical surface of the epithelium every 30 seconds for 10 minutes. The multiple position acquisition feature was used to image many ovarioles in a single imaging session. Cells were segmented and tracked, and tissues were aligned to center, as described above. For stage 6-7 egg chambers, migration rate was calculated based on cell speeds at the central region of the egg chamber between the two poles, to account for variation across the AP axis. For stage 5 and under, all cells were used in the analysis. For younger egg chambers (stage 2-3), only egg chambers in which at least 3 cells could be tracked were included in the analysis. Migration rates were determined from centroid displacement [36]. See also ‘Code’ section below. To quantify migration rates at stage 1 (Fig. 1), we generated kymographs from timelapse movies by drawing single lines through the epithelium in the direction of migration. The migration rate was determined by measuring the slope of 3 kymograph lines and averaging the values. See Barlan et al., 2017 [32] for in-depth methods.

#### Analysis of relative cell movement at basal versus apical surfaces

For each egg chamber, a 30-minute timelapse movie (15 second intervals) was acquired at the basal surface of the epithelium. After 30 minutes, the z-plane was shifted apically and another 30-minute timelapse movie was collected. Cells were segmented and tracked, and centroid positions were obtained as described above. For analysis, cells present for less than 10 consecutive frames were discarded. First, we determined the difference between the first and last position of each cell centroid in all tissues (Fig. S2B). To compare cell movement at basal versus apical surfaces in one egg chamber, we calculated the average cell displacement by finding the total of all cell displacements at each z-plane divided by the number of cells analyzed in that plane (Fig. 4B, S2B).

#### Quantifying polar order from cell centroid tracks

For each egg chamber, a 30-minute timelapse movie was acquired at the basal surface of the epithelium at 15 second intervals. Cells were segmented and tracked, and centroid positions were obtained as described above. Cells present for less than 10 consecutive frames and cells at the boundary of the tissue were not analyzed. The position of cell centroids were defined as 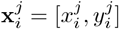 for each cell *j* at time *t* = *i*, where *t* is computed every minute for 30 minutes.

We quantify the centroid movement for cell *j* at time *i* as 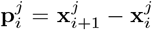. To quantify the coordination between the centroid movements of all cells in the field of view (FOV) over a span of 30 minutes, we compute a polar order parameter *α* for an egg chamber. We calculate *α* from 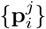 by

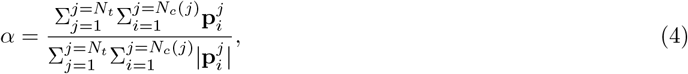

where *N*_*t*_ is the total number of time points and *N*_*c*_(*t*) is the total number of follicle cell centroids in the FOV at both *t* = *i* and *t* = *i* + 1. For perfectly ordered centroid movements where all the cells align in a single direction *α* = 1. For random isotropic centroid movements *α →* 0. See also the ‘Code Availability’ section below.

### Analysis of fixed tissue data

#### Fat2-3xGFP fluorescence intensity quantification

To measure Fat2 brightness at the basal cell edges (Fig. 2B), a single optical section was captured and the Fiji oval selection tool was used to measure the mean intensity of Fat2-3xGFP at the whole basal surface and three cell interiors at the basal surface of each sample. To control for variation between samples, the average of the basal surface cell interiors intensity measurement was subtracted from the total basal surface intensity measurement.

#### Fat2 enrichment at leading-trailing cell edges

As a measure of Fat2 planar polarity at the basal surface, we determined the ratio of Fat2-3xGFP fluorescence intensity at horizontal (migratory leading/trailing) to vertical (lateral) cell edges. To do this, cells were segmented based on a cell membrane marker (anti-DN cadherin) using the pretrained ‘cytoplasm’ model in Cellpose [56]. Next, we segmented the cell-cell edges which are the cellular interfaces between each pair of neighboring cells. Then we measured the angle of each cell-cell edge relative to the egg chamber’s AP axis and Fat2 fluorescence intensity along those edges. Finally, we calculated the average fluorescence intensities of edges with angles between 0° and 10° (migratory leading/trailing) and between 80° and 90° (lateral) and determined the ratio between these values. A detained description of cell-cell edge segmentation can be found in Williams et al., 2022 [33]. See also ‘Code’ section below.

### Reproducibility and statistical analysis

Egg chambers damaged during sample preparation were excluded from analysis. All experiments were performed at least twice. Each replicate included egg chambers pooled from multiple females. Experiments were not randomized, nor was the data analysis performed blind. No statistical method was used to determine sample size. The number of biological replicates (n), statistical tests, and significance can be found in figures or figure legends. All statistical tests were performed in GraphPad Prism10. All data in which statistical tests were performed was tested for normality, and found to follow an approximate normal distribution. Paired t-tests were used for comparison of relative cell movements at basal versus apical surfaces within epithelia (Fig. 4B). A two-way ANOVA with Tukey’s multiple comparisons test was performed on datasets measuring migration rates or Fat2 planar polarity at multiple stages for multiple conditions (Fig. 2B,D, Fig. 3B, S1F). An ordinary one-way ANOVA was performed when two or more conditions were compared within one stage (Fig. S2B).

### Movie generation

Fiji was used to add labels and annotations and export timelapse videos as.avi, which were then converted to.mp4 using HandBrake. Simulations were generated using MATLAB R2023a.

## Acknowledgements

We are grateful to the members of the Horne-Badovinac and Serra labs and to Noah Mitchell for helpful discussions and comments on the manuscript, and to the Bloomington Drosophila Stock Center and FlyBase for the essential reagents and resources they provided for this work. M.S. and S.H.B. further thank The Company of Biologists and the organizers of the 2022 workshop ‘From Physics to Function’ (Buxted Park, UK), where this project was initiated.

## Author Contributions

Conceptualization: Schwabach, Santhosh, Cetera, Serra, Horne-Badovinac; Software: Santhosh, Williams; Experimental data acquisition: Schwabach; Numerical simulations: Santhosh; Formal analysis: Schwabach, Santhosh; Writing: Schwabach, Santhosh, Serra, Horne-Badovinac; Supervision: Serra, Horne-Badovinac; Funding acquisition: Serra, Horne-Badovinac.

## Code Availability

The code used for the experimental analysis and model simulation is available in https://github.com/SreejithSanthosh/egg-chamber-rotation.

## S1 Mechanochemical model of delayed migration egg chamber

**Figure S1.**
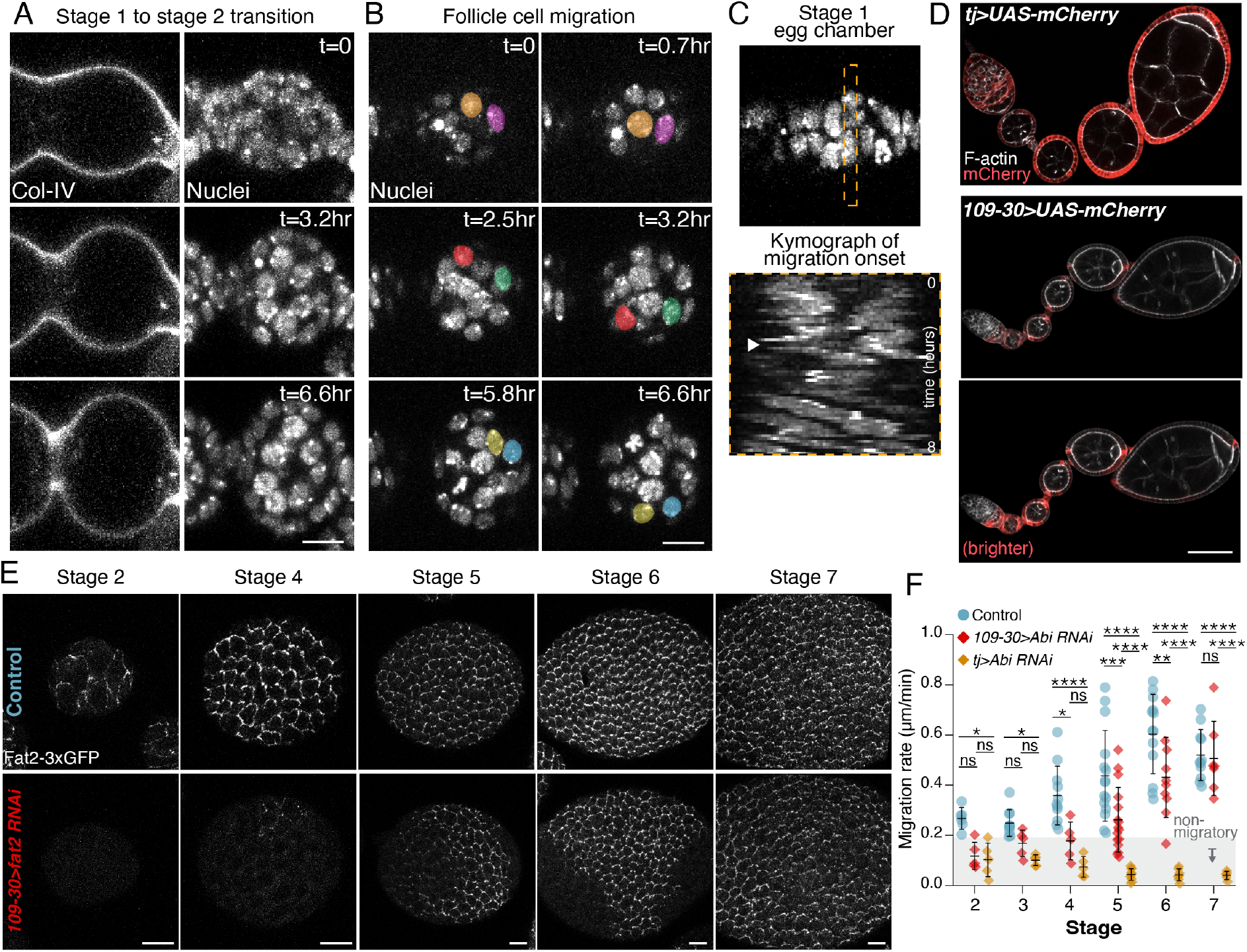
Additional characterization of the onset of migration at stage 1, and delayed-migration onset, related to Figure 1 and 2. **A)** Additional example of stage 1 to stage 2 transition. Movie stills of a transverse section through a stage 1 egg chamber. **B)** Movie stills focused on the follicular epithelium of the same egg chamber as in A. Two different cells are pseudocolored in each row of images to show cell movement. **C)** Additional example of migration onset (arrowhead) at stage 1. Scale bars, 10µm. **D)** Representative images showing the two Gal4 drivers used in this study driving *UAS-mCherry*. The *tj-Gal4* driver is expressed in the follicle cells throughout the stages when the egg chamber rotates, whereas the *109-30-Gal4* driver is primarily expressed in the earliest rotation stages. The first two images were taken with the same microscope settings. The fluorescence signal in the third image has been enhanced to better show the expression pattern. Scale bar, 50µm. **E)** Representative images of Fat2-3xGFP at the basal epithelial surface. *109-30>fat2-RNAi* eliminates Fat2 expression through stage 2, but Fat2 gradually reaches full expression by stage 7 as the RNAi stops being expressed. Scale bar, 10µm. **F)** Quantification of follicle cell migration rates over developmental time. *109-30>Abi RNAi* blocks migration at early stages but migration is indistinguishable from controls by stage 7. Each data point represents one egg chamber. Control values are repeated from Figure 2D. Two-way ANOVA with Tukey’s multiple comparisons test; ns, *p >* 0.05, \**p <* 0.05, \*\**p <* 0.01, \*\*\**p <* 0.001, \*\*\*\**p <* 0.0001. In order on graph, *n* = 6, 5, 5, 10, 5, 5, 11, 5, 5, 15, 16, 10, 13, 10, 7, 11, 7, 5. Bars represent mean ± SD.

**Figure S2.**
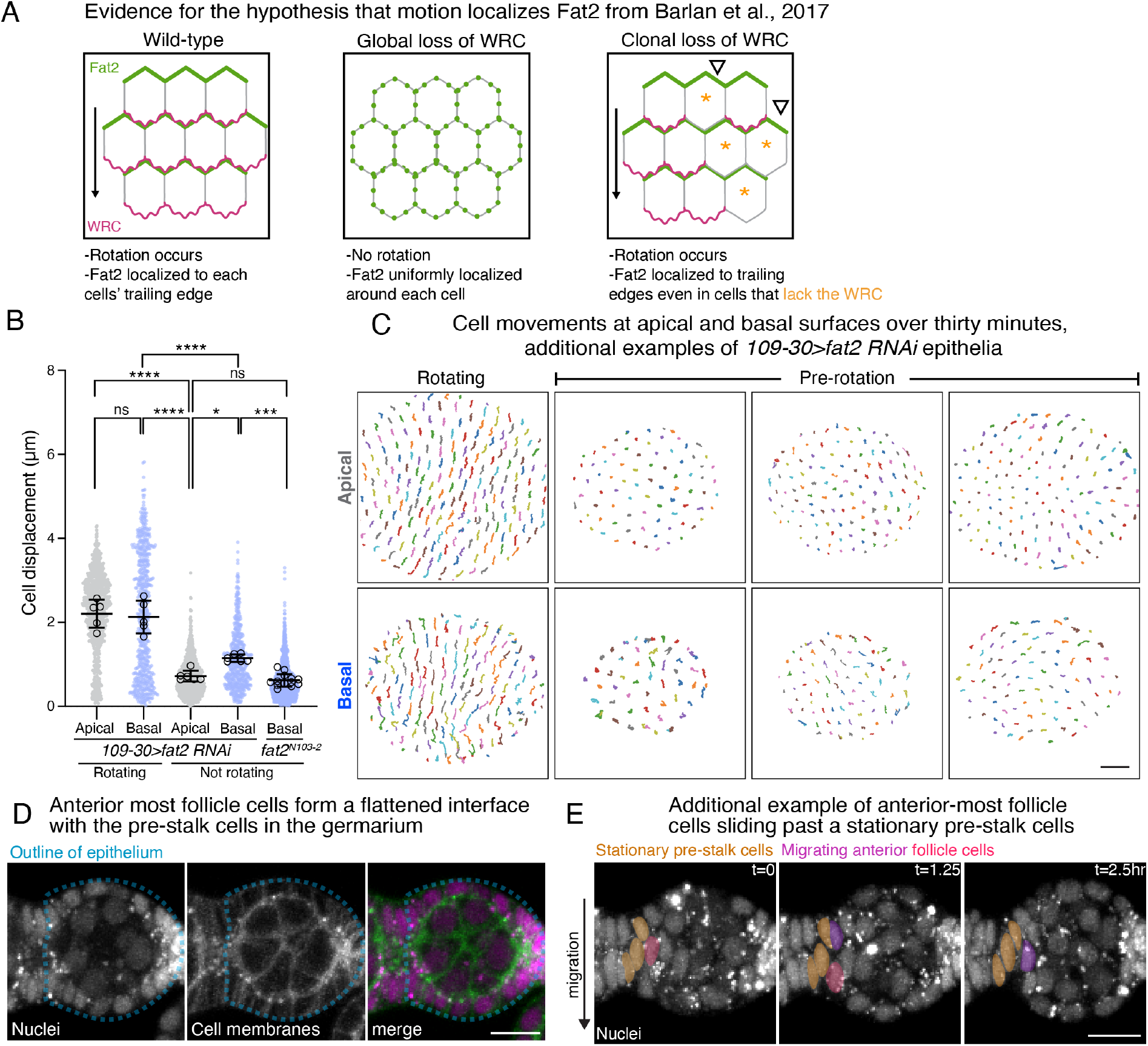
Relationship between Fat2 polarity and global rotation; additional characterization of cell movement at apical and basal surface planes; additional example of follicle cell/pre-stalk interaction. Related to Figures 3, 4 and 6. **A)** Illustration of the experiment from Barlan et al., 2017 that first suggested that the localization of Fat2 to the trailing edge of each follicle cells depends on tissue motion. **B)** Quantification of cell displacement at different imaging planes. Closed dots represent individual cells across multiple egg chambers, black open circles represent the average value for each egg chamber. For egg chamber averages: Ordinary one-way ANOVA with Tukey’s multiple comparisons test; ns, *p >* 0.05, \**p <* 0.05, \*\*\**p <* 0.001, \*\*\*\**p <* 0.0001. In order on graph, *n* = 5, 5, 6, 6, 13. Bars represent mean ± SD. **C)** Additional examples of cell centroid tracks over 30 minutes at apical and basal surface planes in rotating or not-rotating *109-30>fat2-RNAi* egg chambers. **D)** Transverse section of the stage 1 egg chamber shown in Fig. 6B, used to generate the diagram in Fig. 6A, highlighting the shape of the follicular epithelium in a rotating egg chamber at stage 1. Cell membranes were visualized using CellMask. **E)** Movie stills of maximum intensity projections of nuclei generated from a 2.5 hour movie. Stationary pre-stalk cell nuclei are pseudocolored gold, rotating pre-polar and follicle cell nuclei are pseudocolored magenta and blue, respectively. Scale bars = 10µm

### S1.1 Mechanics of the egg chamber

We model the egg chamber as a rigid body, as there are negligible neighbor exchanges between follicle cells, and these cells tightly adhere to the germ cells. Egg chambers have an ellipsoidal geometry and are slightly elongated along the AP axis at stages 5-6, when the delayed onset of rotation occurs. We parametrize the egg chamber (i.e. a surface *M*, Fig S3A) as

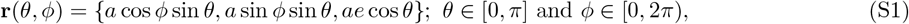

where *a* is the radius of the equatorial cross-section and *e* is the eccentricity (Fig. S3B). At each point on the egg chamber **r**(*θ, ϕ*), the tangent vector basis (_1_*ζ*, _2_*ζ*) and the surface element **dA** (Fig. S3A) are given by

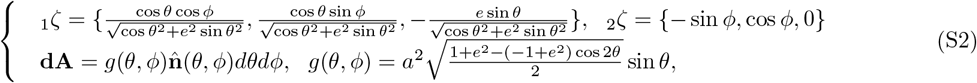

where 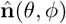 is the outward normal vector. To describe the dynamics, we define an inertial (i.e. fixed) frame 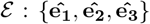 and a body-frame 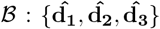 that rotates with the egg chamber and moves with its center of mass, with 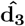 oriented along the long axis (Fig. S3B). The motion of a rigid body in 3D can be described by its center of mass velocity **v** and its angular velocity *ω*. The dynamics of the egg chamber given external forces and torques obeys linear momentum and angular momentum principles in the over-damped limit, which in the body-frame *B* [1], read

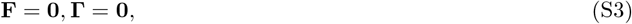

where **F, Γ** are the total applied force and torque, which we model below.

**Figure S3.**
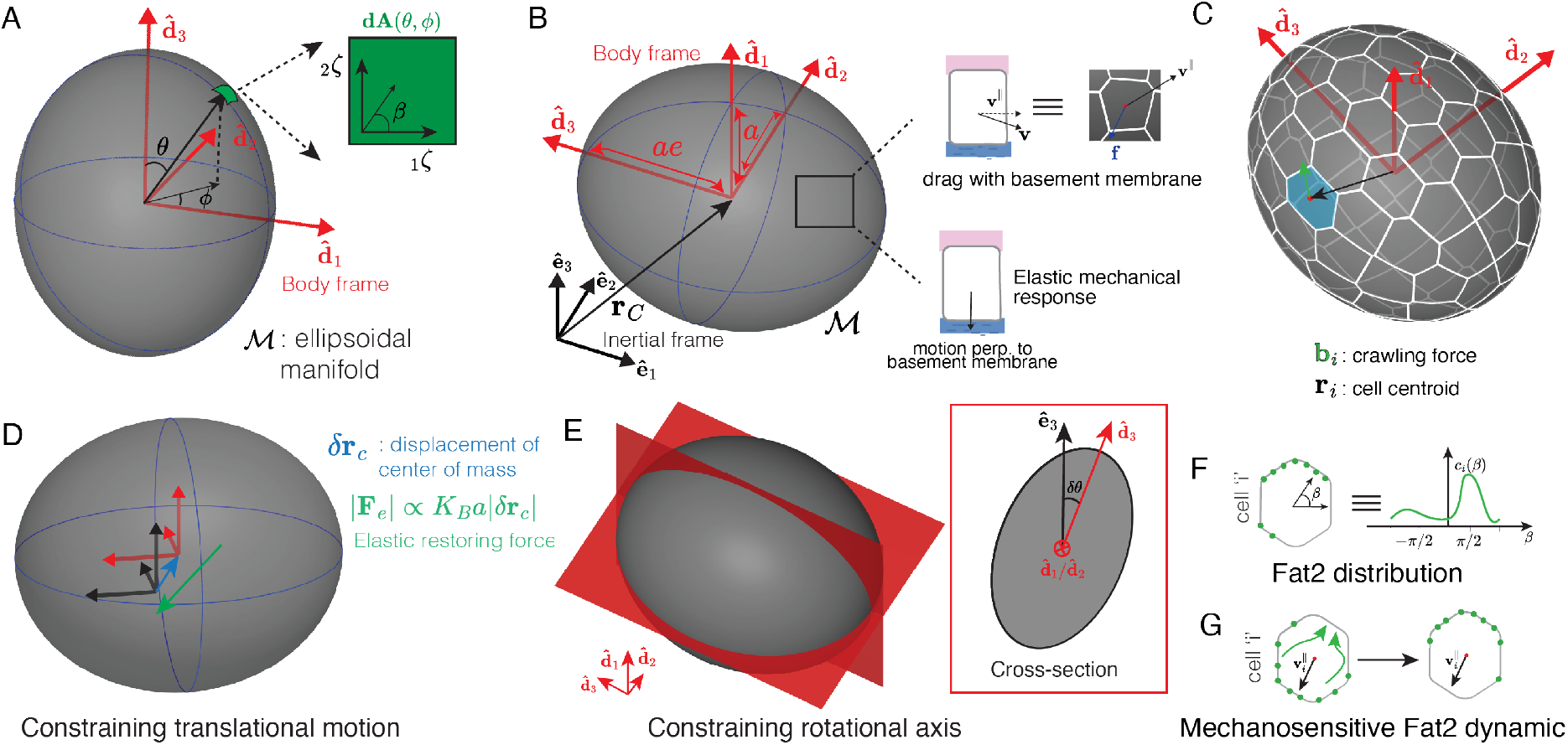
Biophysical model of egg chambers during delayed migration onset. (a) Egg chamber geometry is described using an ellipsoidal manifold *ℳ* parameterized using angles (*θ, ϕ*) in the body frame 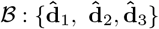. The area element **dA** at (*θ, ϕ*) has tangent bases vectors *{*_1_*ζ*, _2_*ζ}* and *β* is the angular coordinate on the tangent plane. (b) The dynamics can be described both in the inertial frame *ℰ* and the body frame *ℬ*. The basement membrane provides viscous (elastic) resistance force to motion tangential (perpendicular) to it. (c) Follicle cell positions described in the body frame. (d-e) Elastic response of the basement membrane to motion of the egg chamber normal to the interface constrains translational motion (d) and rotational axis (e). (f) The Fat2 concentration for cell *i* is described using *c*^*i*^(*β*). (g) The Fat2 concentration is mechanosensitive.

#### Tangential drag force at the basement membrane

The basement membrane encases the egg chamber and resists its relative tangential motion via viscous drag due to remodeling of integrin-based adhesion complexes (Fig S3B). The basement membrane responds elastically to motion normal to the surface, which we model and discuss the consequences in Sec. S1.1.1. We model the tangential surface drag force density as

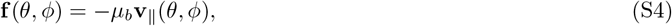

where *µ*_*b*_ is the viscous drag coefficient and **v**_*∥*_ is the component of the velocity tangential to the surface at (*θ, ϕ*).

The velocity **v**(*θ, ϕ*) of a surface element is given by

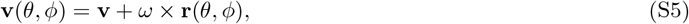

where **r**(*θ, ϕ*) is the position vector of the surface element with respect to the center of mass of the egg chamber. The total force and torque generated by the local viscous drag forces given in Eq. (S4) are obtained by integrating over the entire surface of the egg chamber (Eq. (S1)). Given that *e ≈* 1.2 for a stage 5 egg chamber we compute the total viscous friction force **F**_**f**_ and torque **Γ**_**f**_ (expressed in *B*) by expanding about *e* = 1 in linear order, which gives

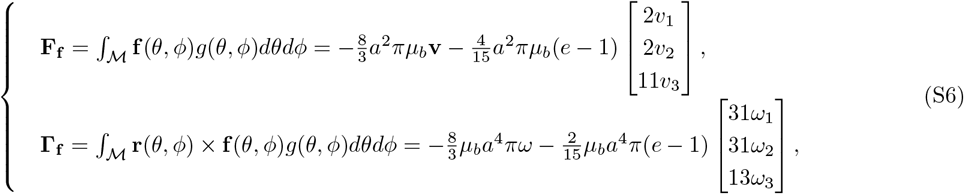

where **v** = [*v*_1_, *v*_2_, *v*_3_] and *ω* = [*ω*_1_, *ω*_2_, *ω*_3_] are expressed in the body-frame *ℬ*.

#### Protrusive crawling force

The follicle cells use cryptic lamellipodia at their basal surfaces to crawl on the basement membrane [2]. We model these active point forces **b**^**i**^ acting at the cell-centroid **r**^**i**^ *∈ M* (Fig S3C) on the basal surface of cell *i*. Therefore, the total crawling (or active) force **F**_*a*_ and torque **Γ**_*a*_ exerted by the basal surface protrusions are given in the body frame by

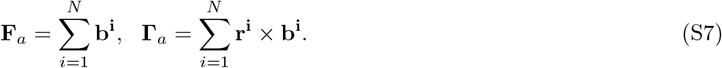

#### S1.1.1 Elastic confinement due to basement membrane

The basement membrane elastically resists motion perpendicular to the egg chamber-basement membrane interface.

The elastic surface force density is given by

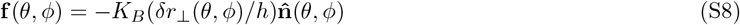

where *K*_*B*_, *h* are the stiffness and thickness of the basement membrane, and *δr*_*⊥*_(*θ, ϕ*) is the displacement of the area element normal to the interface. For an egg chamber with the center of mass displaced by 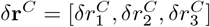 in the body-frame *ℬ* and body-frame orientation given by rotation matrix **A**,

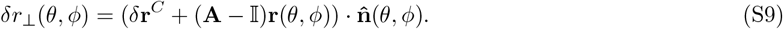

The rotation matrix **A** gives the orientation of the body-frame with respect to the inertial-frame. The orientational rest-configuration of the egg chamber is assumed to be **A** = 𝕀 and we analyze small deviations *δθ* of the egg chamber AP axis away from 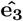 (Fig S3D-E). Integrating over the egg chamber, we obtain the total force and torque to linear order in *e* represented in the *ℬ* frame:

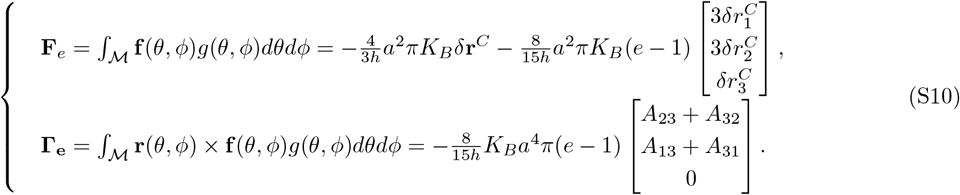

Active crawling forces drive the rigid rotation. An order of magnitude estimate of the active force **F**_*a*_ and torque **Γ**_*a*_ is given by

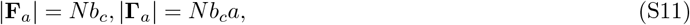

where *b*_*c*_ is the characteristic crawling force of one cell. *b*_*c*_ = *σ*_*a*_*A*_*c*_, where *σ*_*a*_ is the mean traction stress generated by a crawling cell and *A*_*c*_ is the cell area. There are currently no available experimental measurements of *σ*_*a*_ in the *Drosophila* egg chamber. Instead, we use an estimate of traction stresses *≈* 10 Pa measured from collectively migrating epithelial cells in another *in vivo* system [3]. Equating the magnitude of active crawling and elastic response force (torque), we can estimate the theoretical maximum translation (rotational motion) of the egg chamber. Using the values of *K*_*B*_ *≈* 50kPa, *h ≈* 90*nm, a ≈* 25*µm, e ≈* 1.2 [4–7] we get,

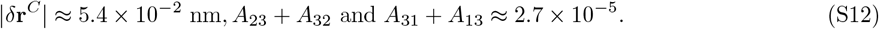

Therefore, since the motion of the center of mass is significantly smaller than the dimensions of the egg chamber (|*δ***r**^*C*^| *<< a*), we assume that the center of mass remains stationary for all our simulations. The entries *A*_23_, *A*_32_, *A*_31_, *A*_13_ of the orientation matrix **A** quantifies the deviation of the AP axis (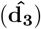) from the rest orientation (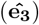). Since Eq. (S12) holds for any rotation of the egg chamber about an axis in the 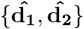plane, we can independently infer that *A*_32_, *A*_23_, *A*_31_, *A*_13_ *≈* 2.7 *×* 10^*−*5^. This demonstrates that rotation of the egg chamber is confined to its long axis, consistent with experimental observations [8]. This constraint on rotational axis does not hold for spherical egg chambers (*e →* 1), which can rotate along any axis (See Sec. S1.5).

### S1.2 Crawling force and cell polarity dynamics

#### S1.2.1 Fat2 polarity dynamics

We model the Fat2 distribution for cell *i* as *c*^*i*^(*β, t*), where *β* is the angular coordinate in the tangent space of *ℳ* at the cell-centroid position **r**^*i*^ (Fig. S3F). Previous work [9] has demonstrated that Fat2 localizes to cell’s trailing edges, and also suggests that tissue rotation may play a role in localizing Fat2. Therefore, we construct a minimal model of mechanosensitive Fat2 dynamics as

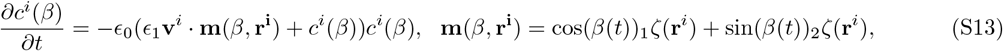

where **v**^*i*^ = **v** + *ω ×* **r**^*i*^ is velocity of cell *i, ϵ*_0_ is a characteristic rate constant for the dynamics of Fat2 polarity and *ϵ*_1_ sets the strength of coupling between migration and Fat2 polarity. Intuitively, Eq. (S13) models an increase in Fat2 polarization to the cell edge in the direction opposite of motion (Fig. S3G).

#### S1.2.2 Crawling force dynamics

Fat2 acts at cells’ trailing edges to create a stable protrusive domain in the leading edge of the neighboring cell behind [9]. Thus we assume the direction of the crawling force **b**^*i*^ to be influenced by Fat2 polarity. For simplicity, we model the activity of Fat2 as cell-autonomous and the crawling force **b**^*i*^ as

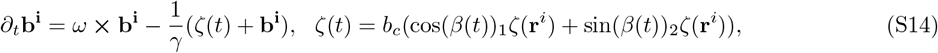

where *β*(*t*) is sampled from the probability distribution *P*_*i*_(*β*) *∝ c*_*i*_(*β*), *b*_*c*_ is the characteristic scale of the force generated by cell protrusions and *γ* is the timescale taken by the protrusion to respond to changes in Fat2 polarity encoded inζ(*t*). The term *ω* ***×* b**_**i**_ arises from differentiation in rotating frames and maintains the consistency of the protrusion direction during rigid rotation of the egg chamber.

### S1.3 Nondimensional equations of motion and parameter analysis

Equations (S6-S7,S13-S14) constitute a mechanochemical model of the egg chamber. We nondimensionalize the variables using the transformations *t → t*_*c*_*t, ω → ω*_*c*_*ω*, **r**_*i*_ *→ a***r**_*i*_, **v** *→ ω*_*c*_*a***v, b**_**i**_ *→ b*_*c*_**b**_*i*_, *c*_*i*_ *→ c*_*c*_*c*_*i*_ andζ *→ b*_*c*_*ζ*; where *t*_*c*_, *ω*_*c*_,*v*_*c*_, *b*_*c*_, *c*_*c*_ are the characteristic time-scale of rotation initiation, angular-speed of egg chamber, force scale of protrusions and characteristic concentration scale of Fat2. Rotation initiation occurs over a time-scale *t*_*c*_ *≈* 1*hour*. The radius of the egg chamber equatorial cross-section *a ≈* 25*µm* during rotation initiation. The characteristic angular velocity 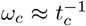 for an egg chamber during delayed rotation initiation. We further use the fact that after symmetry breaking, the egg chamber rotates in a single direction with an angular velocity *ω ω*_*c*_. In this regime, torque from crawling forces must balance torque from drag with the basement membrane, which, using the angular momentum balance, gives *b*_*c*_*aN≈ µ*_*b*_*a*^4^*ω*_*c*_ (order of magnitude calculations), leading to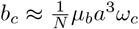. Due to the strong elastic confinement from the basement membrane, the center of mass remains stationary and rotations are restricted to the long axis (See Sec. S1.1.1). Therefore, we obtain the non-dimensional equations of motion

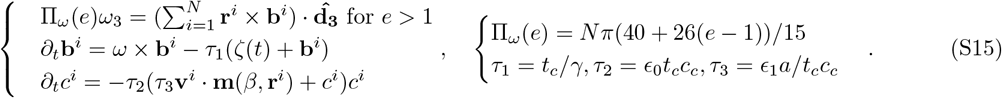

For simulations of the delayed onset of rotation, we use *e* = 1.2 and *N* = 300 consistent with experimental observations [7]. To compute the cell centroid locations **r**^*i*^ for a given number of follicle cells *N*, we initialized it by placing *N* equi-distant points on the surface of an ellipsoid.

#### S1.3.1 Parameters *τ*_1_, *τ*_2_ **and** *τ*_3_

Rotation initiation occurs over a time-scale *t*_*c*_ *≈* 1*hour*, and the time-scale over which protrusions respond to changes in Fat2 distribution is *γ ≈* 3 min [10]. Therefore, we set *τ*_1_ = 20. There is currently no reliable experimental estimate for (*τ*_2_, *τ*_3_). Therefore, we numerically explore the parameter space and observe that the egg chamber undergoes symmetry breaking for all positive values, but the time taken to break symmetry changes. We quantify this difference by estimating the time taken to reach 95 percent of the ensemble average of the steady state angular velocity |*ω*_3_|^*s*^ after symmetry breaking (See SI Fig S4 A) for a parameter set. Increasing (*τ*_2_, *τ*_3_) decreases the time taken for the egg chamber to symmetry break (See SI Fig S4 B-C).

**Figure S4.**
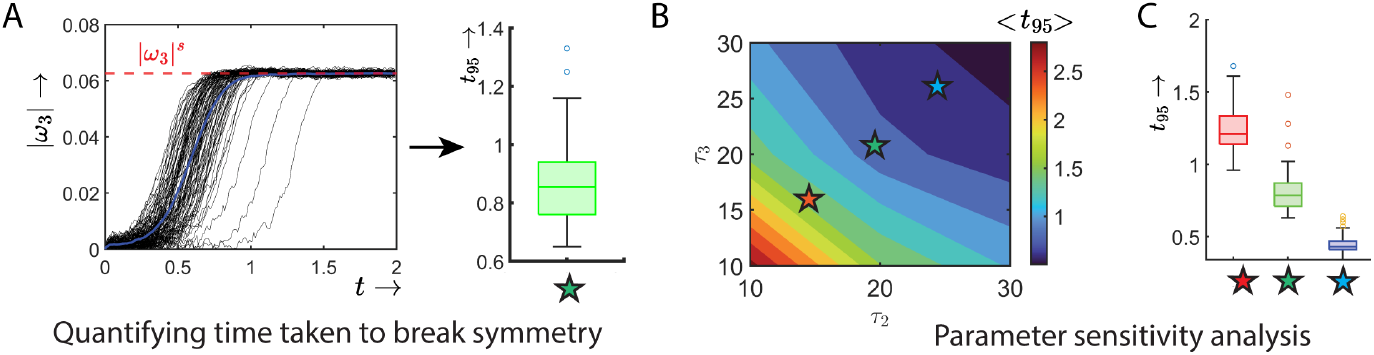
Sensitivity analysis to changes in. (*τ*_2_, *τ*_3_). (a) *N* = 100 runs of the theoretical model of a delayed egg chamber for the parameter values (*τ*_2_, *τ*_3_) = (20, 20). (b-c) The ensemble average time taken *< t*_95_ *>* to reach 95 percent of the steady-state symmetry broken angular velocity |*ω*_3_|^*s*^ for different parameter sets.

### S1.4 Numerical implementation

We simulate an egg chamber undergoing delayed rotation initiation by solving Eqns. (S15) using forward Euler integration method with time-step *dt* = 10^*−*2^ in MATLAB. The evolution of the body frame is tracked using a rotation matrix **A**(*t*) which relates vectors in the inertial-frame *ℰ* and body-frame *ℬ* as

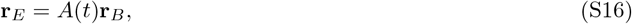

where **r**_*E*_, **r**_*B*_ are the coordinates of the same vector in the inertial and body frames. The rotation matrix **A**(*t*) evolves using the relation

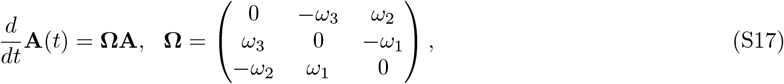

where *ω* = [*ω*_1_, *ω*_2_, *ω*_3_] is the angular velocity of the egg chamber represented in the inertial frame. For an ellipsoidal (*e >* 1) egg chamber *ω*_2_, *ω*_1_ = 0 due to the strong confinement from the basement membrane (Sec S1.1.1).

### S1.5 Spherical egg chamber

In contrast to an elongated egg chamber, the elastic confinement from the basement membrane does not constrain the rotational axis in a spherical egg chamber (Sec S1.1.1 for details). Therefore, using the same non-dimensionalization scheme as introduced in Sec. S1.3 the non-dimensional equation of motion for a spherical egg chamber is given by

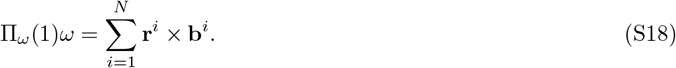

To simulate rotation initiation in a spherical egg chamber, we use Eq. (S18) for the rotational dynamics and the same equations for Fat2 concentration and crawling force as in the elongated egg chamber (Eq. (S15)). We simulate these equations with the same parameters *τ*_1*−*3_ as the delayed migration egg chamber, but with *N* = 100, as quantified for the stage 1 egg chamber. Starting from an isotropic initial condition (*c*^*i*^(*β*, 0) = 1*/*2*π*, **b**^*i*^(0) = **0**), the spherical egg chamber undergoes symmetry breaking (Fig S5A-C) but with a randomly oriented rotational axis (Fig S5D).

**Figure S5.**
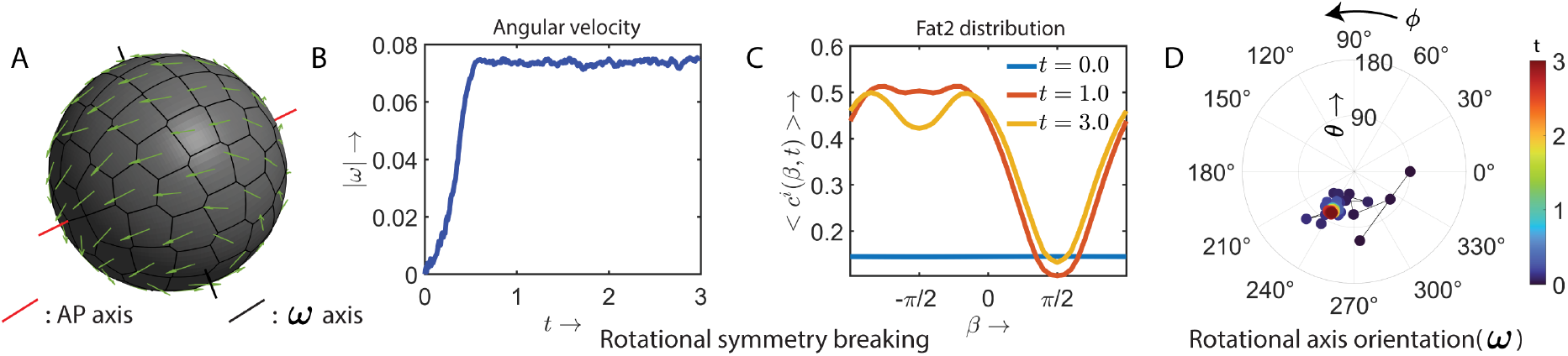
Random rotational axis in spherical egg chambers. (a) Rotation of a spherical egg chamber. (b-c) The egg chamber undergoes symmetry breaking, generating sustained rotations. (d) The orientation of the angular velocity *ω* described in the inertial frame *ℰ* is quantified using spherical coordinates (*θ, ϕ*). The time-evolution of the quantities in a-d is given in Movie 10. We mark the future AP axis (red) for reference.

## S2 Mechanochemical model of stage 1 egg chamber

### S2.1 Mechanics of stage 1 egg chambers

The mechanics of stage 1 egg chambers is distinct from the later stages due to the contact of follicle cells with the pre-stalk cells (Fig. S6A). We model the mechanical interaction with the stationary pre-stalk cells to elucidate how the stage 1 egg chamber undergoes symmetry breaking. The stage 1 egg chamber is spherical (*e* = 1), but with a flat contact with the pre-stalk cells. The spatial extent of the contact region is quantified using the polar angle *θ*_*s*_. We use a similar framework to the model introduced in Sec. S1.1, but accounting for the interaction with the pre-stalk cells. The strong confinement from the basement membrane, discussed in Sec. S1.1.1, holds even for the stage 1 egg chamber; therefore, the center of mass is fixed. We next discuss the changes in the external torques of this system compared to the delayed migration model.

#### Tangential drag force at the basement membrane

The tangential drag force is the same as for the egg chamber at the time of delayed-rotation onset Eq. (S4), but the surface of integration is given by *ℳ*_*b*_ (Fig. S6B):

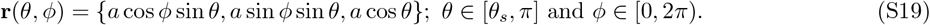

The total (friction) torque due to the drag with the basement membrane is given by

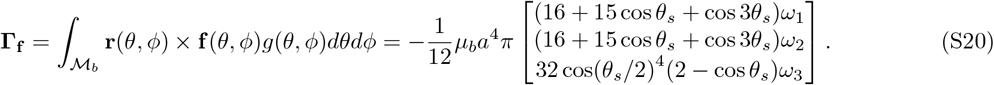

#### Tangential drag force at the pre-stalk interface

We assume a viscous drag between the stationary pre-stalk and the egg chamber for motion tangential to the interface. The surface drag force density is given by

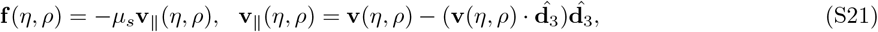

where (*η, ρ*) is the polar parameterization of the pre-stalk interface (See Fig S6B), *µ*_*s*_ is the drag coefficient for the viscous interaction between the egg chamber and the pre-stalk, and **v**(*η, ρ*) is the velocity of the area element. The torque exerted by the viscous force given in Eq. (S21) is given by

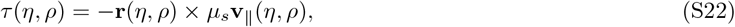

where **r**(*η, ρ*) is the position vector of the area element at (*η, ρ*). Integrating Eq. (S22) over the egg chamber/pre-stalk interface *ℳ* _*s*_, we get the corresponding total torque exerted by this viscous force:

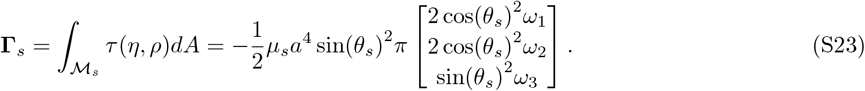

#### Elastic confinement at the pre-stalk interface

Motion of an area element **dA** at (*η, ρ*) on the stalk-egg chamber interface normal to it generates an elastic resisting force. The elastic surface force density is given by

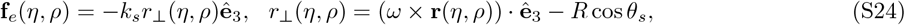

where *k*_*s*_ is the stiffness of the pre-stalk cells and *r*_*⊥*_(*η, ρ*) is the displacement of the area element normal to the interface (See Fig. S6C). Similar to the tangential drag force given above, we compute the torque and integrate over the interface *ℳ* _*s*_ to get the total elastic torque

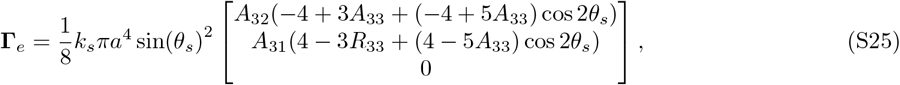

where *A*_*ij*_ are the components of the rotation matrix that transforms vectors from the body frame *ℬ* to *ℰ*.

**Figure S6.**
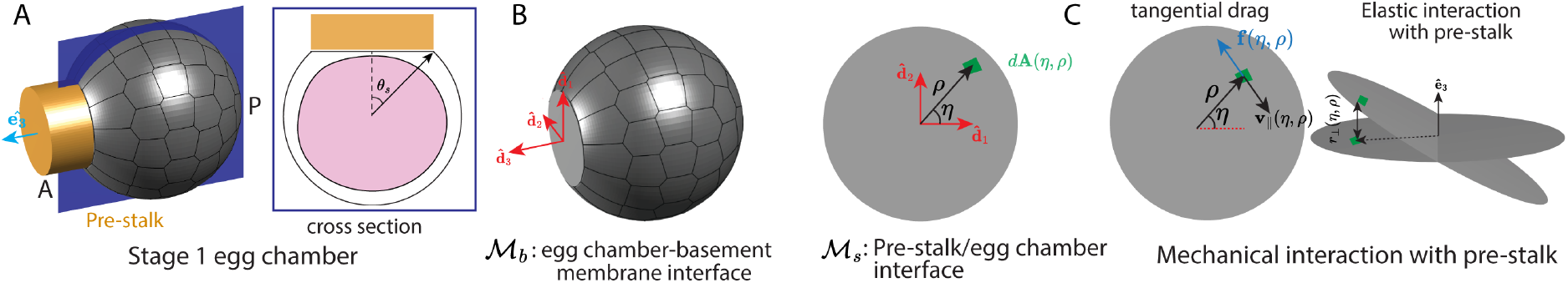
Model of mechanical interactions in a stage 1 egg chamber. (a) Geometry of a stage 1 egg chamber. (b) The pre-stalk/egg chamber interface *ℳ* _*s*_ is parameterized by the polar coordinate (*η, ρ*). (c) Motion tangential (normal) to the interface is assumed to have a viscous (elastic) resisting force.

#### S2.1.1 Non-dimensional equation of motion and parameter selection

We use the same non-dimensionalization scheme in Sec. S1.3. The Fat2 polarity and protrusion dynamics are as above, while the angular momentum balance is different, given by

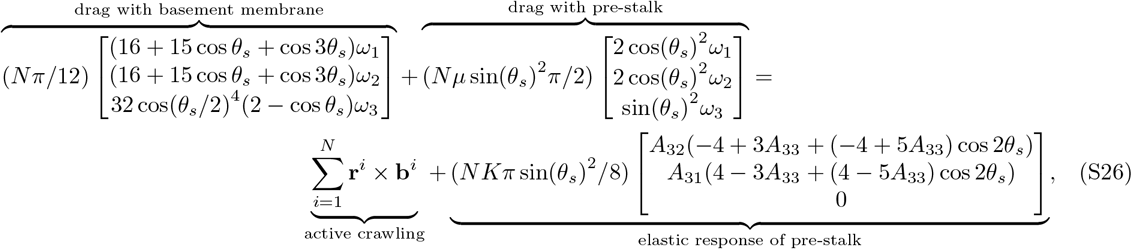

where *µ* = *µ*_*s*_*/µ*_*b*_ and *K* = *k*_*s*_*/µ*_*b*_*ω*_*c*_. To simulate these equations, we use the same parameter values for *τ*_1*−*3_ as the delayed migration egg chamber, but with *N* = 100 corresponding to a stage 1 egg chamber. There are no experimental measurements of either *µ* or *K*; therefore, we set *µ* = 1 and *K* = 20. We observe no significant change in the qualitative behavior of the model even for significant changes in these parameter values, consistently generating egg chamber rotations along the pre-stalk axis at steady state.

## Movie legends

**Movie 1. Egg chamber rotation occurs within the germarium** Related to Figure 1. An example egg chamber transitioning from stage 1 to stage 2, which is signified by the basement membrane (Col-IV-GFP) becoming continuous over the anterior portion of the egg chamber. This egg chamber undergoes sustained rotations for the length of the movie. This sample was used to generate Fig. 1E,F. Nuclei were visualized using SpyDNA, which labels all follicle cell and germ cell nuclei. Time stamp shows hours:minutes.

**Movie 2. Egg chamber rotation initiates during stage 1**. Related to Figure 1. The follicular epithelium of two different stage 1 egg chambers undergoing either sustained rotation (left) or initiating rotation (right). The yellow arrow appears at the onset of rotation. These egg chambers were used to generate the kymographs in Fig. 1I. Nuclei were visualized using SpyDNA, which labels all follicle cell and germ cell nuclei. Time stamp shows hours:minutes.

**Movie 3. *109-30****>****fat2 RNAi* delays the onset of migration**. Related to Figure 2. A control ovariole (left) and *109-30>fat2* RNAi ovariole (right) each containing a stage 3 and stage 6 egg chamber, focused on the follicular epithelium. Temporal knock down of Fat2 using the 109-30-Gal4 driver (right) blocks migration specifically in the stage 3 egg chamber, while migration occurs normally in the adjacent stage 6 egg chamber. Follicle cells in both stage 3 and 6 egg chambers migrate in the control ovariole (left). Cell membranes were visualized using CellMask.

**Movie 4. Rotation can initiate *ex vivo*, after a delay in *fat2* expression**. Related to Figure 2. The follicular epithelium of (left) two control egg chambers undergoing continuous rotation *ex vivo*, and (right) four examples of *109-30>fat2 RNAi* egg chambers undergoing delayed onset of rotation *ex vivo*. All examples are of stage 5 egg chambers. Indy-GFP labels follicle cell membranes. Time stamp shows hours:minutes.

**Movie 5. Tracking of cell centroid positions**. Related to Figure 4. The basal surface of a *109-30>fat2 RNAi* epithelium in a stage 5 egg chamber pre-rotation to demonstrate how cell centroid positions are used to track cell movement. Cell membranes were visualized using CellMask.

**Movie 6. Follicle cells’ basal surfaces are highly dynamic before rotation begins**. Related to Figure 4. Example of a *109-30>fat2 RNAi* epithelium that is rotating, a *109-30>fat2 RNAi* epithelium pre-rotation, and a *fat2*^*N103-2*^ epithelium that will never rotate. In rotating egg chambers, the apical and basal surfaces move in concert, while motion in pre-rotating egg chambers is restricted to the basal epithelial surface. This local motility is dependent on Fat2 activity. All examples are of stage 5 egg chambers. Samples were used to generate Fig. 4A. Cell membranes were visualized using CellMask.

**Movie 7. Simulation of the delayed onset of egg chamber rotation**. Related to Figure 5, panel F. The time evolution of the biophysical model of a delayed migration egg chamber with an ellipsoidal geometry.

**Movie 8. Anterior-most follicle cells migrate past pre-stalk cells in stage 1 egg chambers**. Related to Figure 6. Timelapse of a stage 1 egg chamber with pre-stalk cells highlighted in orange and anterior-most follicle cell in magenta. Over time, the anterior-most follicle cells slide past the pre-stalk cells which remain stationary. This sample was used to generate Fig. 6B. Nuclei visualized with ubi-mRFP-NLS. Time stamp shows hours:minutes.

**Movie 9. Simulation of the onset of rotation at stage 1**. Related to figure 6, panels F and G. The time evolution of the biophysical model of a stage 1 egg chamber.

**Movie 10. Simulation of spherical egg chamber rotation**. Related to figure S5. The time evolution of the biophysical model of a delayed migration egg chamber with a spherical geometry.

